# Spatial information transfer in recurrent place-cell networks depends on excitation-inhibition balance, neural-circuit heterogeneities, and trial-to-trial variability

**DOI:** 10.64898/2026.04.02.716259

**Authors:** Rituparna Roy, Rishikesh Narayanan

## Abstract

A key challenge in understanding spatial navigation and memory is explaining how hippocampal networks sustain robust spatial information transfer despite pronounced trial-to-trial variability and pervasive neural-circuit heterogeneities. Although hippocampal heterogeneities and physiological variability are well-characterized, circuit-scale understanding of stable information transfer in recurrent place-cell networks remains limited. Here, we first show that even recurrent networks composed of intrinsically identical neurons and receiving identical place-field inputs express pronounced neuron-to-neuron variability in spatial tuning profiles, place-field widths, subthreshold ramp amplitudes, and spatial information transfer. Introduction of intrinsic within-type heterogeneities to excitatory and inhibitory neurons further increased diversity in firing properties, but strikingly improved robustness of spatial information transfer under high trial-to-trial variability. Although strengthening inhibition expectedly narrowed place fields and reduced firing across all networks, the impact of inhibition on information-transfer profiles was stronger in heterogeneous networks manifesting high degree of trial-to-trial variability. Across networks, increasing trial-to-trial variability reduced information transfer and shifted the spatial location of peak information from the high-slope regions of the tuning curve to its peak-firing location. Finally, incorporating afferent heterogeneities allowed neurons to be tuned to distinct place-field centers, reducing peak information values while amplifying neuron-to-neuron diversity in information-transfer profiles. Together, we demonstrate that excitation-inhibition balance, trial-to-trial variability, within-type heterogeneities, and afferent input diversity jointly regulate spatial information transfer in recurrent place-cell circuits. We also highlight intrinsic heterogeneities as substrates for enhanced robustness of spatial coding against perturbations. Importantly, the convergence of multiple disparate mechanisms in yielding similar information-transfer profiles underscores degeneracy as a fundamental organizing principle in neural-circuit physiology.

## INTRODUCTION

Biological neural networks are composed of interacting neurons that are tuned to respond to specific stimulus features, with different neurons in the network typically responding to distinct attributes of the stimulus space. For instance, individual neurons in the mammalian primary visual cortical network respond to specific orientations of visual stimuli, with different neurons tuned to respond maximally to distinct orientations (Hubel & Wiesel, 1962). Place cells in the mammalian hippocampal network are tuned to specific spatial locations, with each place cell tuned to distinct spatial locations (Per Andersen, Morris, Amaral, Bliss, & O’Keefe, 2007; E. I. Moser, Kropff, & Moser, 2008; M. B. Moser, Rowland, & Moser, 2015; John O’Keefe, 1976; J. O’Keefe & Dostrovsky, 1971). The selectivity of individual neurons to specific parts of the stimulus space is typically represented by tuning curves that quantify individual neuronal responses to the entire stimulus space. Networks built with neurons that span the entire feature space through their disparate sets of tuning curves have been implicated in precisely and efficiently encoding information about natural stimuli. Relationships between neuronal tuning curves and efficient information transfer have been investigated towards assessment of biological systems from a sensory coding perspective (Attneave, 1954; Barlow, 1961; Bell & Sejnowski, 1997; Brenner, Bialek, & Van Steveninck, 2000; Fairhall, Lewen, Bialek, & de Ruyter van Steveninck, 2001; Laughlin, 1981; Simoncelli, 2003; Simoncelli & Olshausen, 2001) as well as from a single neuron perspective (Andrews & Iglesias, 2007; Lundstrom, Higgs, Spain, & Fairhall, 2008; Narayanan & Johnston, 2012).

The relationship between the tuning curve of neuronal responses and the information transfer profiles has been well explored across different sensory modalities (Butts, 2003; Butts & Goldman, 2006; DeWeese & Meister, 1999; Montgomery & Wehr, 2010). Despite this, the relationship between spatial tuning profiles and spatial information transfer has not been assessed in networks of place cells interacting within the hippocampal circuitry. In one-dimensional arena, place cells respond to distinct locations across the arena through a bell-shaped response profile, with the peak of the curve representing the center of the place field (Buzsaki & Moser, 2013; Dombeck, Harvey, Tian, Looger, & Tank, 2010; Geisler, et al., 2010; Harvey, Collman, Dombeck, & Tank, 2009; M. B. Moser, et al., 2015). Previous studies have assessed the emergence of spatial tuning profiles in a heterogeneous population of hippocampal single neurons (Basak & Narayanan, 2018, 2020; A. Roy & Narayanan, 2021; Seenivasan & Narayanan, 2020), apart from studying the relationship between spatial tuning profile and spatial information transfer in single neuronal models (A. Roy & Narayanan, 2021). However, the dependence of spatial information transfer and spatial tuning curves of neurons in a recurrent network of place cells, such as the CA3 network (P. Andersen, Morris, Amaral, Bliss, & O’Keefe, 2006; Bennett, Gibson, & Robinson, 1994; Le Duigou, Simonnet, Telenczuk, Fricker, & Miles, 2014; Rolls, 2007), on circuit-level and single-neuron parameters has remained surprisingly unexplored. To address these lacunae, in this study, we systematically study the relationship between spatial information transfer and spatial tuning curves of neurons in spiking recurrent networks of place cells endowed with different forms of heterogeneities and different degrees of trial-to-trial variability.

To quantitatively link the place-field firing profile, which is a rate-coded tuning curve of every location in the spatial environment, to the associated spatial information transfer, it is essential that the information metric is available for every location (stimulus value) as well. We used stimulus-specific information (SSI) metric that quantifies the amount of information that the responses of neurons convey about any particular stimulus (Butts, 2003; Butts & Goldman, 2006; DeWeese & Meister, 1999; Montgomery & Wehr, 2010; A. Roy & Narayanan, 2021). As SSI and its relationship to tuning curves have been linked to trial-to-trial variability (Butts, 2003; Butts & Goldman, 2006; DeWeese & Meister, 1999; Montgomery & Wehr, 2010; A. Roy & Narayanan, 2021), we computed SSI for each neuron in the network for different degrees of trial-to-trial variability. As previous studies have emphasized the importance of neural heterogeneities on circuit function (Dahmen, et al., 2025; Gast, Solla, & Kennedy, 2024; Mishra & Narayanan, 2019, 2021b; Mittal & Narayanan, 2021; Saini & Narayanan, 2025; Santhosh & Narayanan, 2025), we assessed the impact of different forms of neural heterogeneities on spatial tuning curves, spatial information transfer, and the relationship between the two. Specifically, we systematically assessed the impact of neuronal intrinsic heterogeneities, synaptic heterogeneities in the form of the balance between excitatory and inhibitory synapses onto individual neurons (Bhatia, Moza, & Bhalla, 2019; Káli & Dayan, 2000; Mackwood, Naumann, & Sprekeler, 2021; Murphy & Miller, 2009; Sprekeler, 2017; Sun, et al., 2017; Van Vreeswijk & Sompolinsky, 1996; Yu, Shen, Wang, & Yu, 2018; Shanglin Zhou & Yuguo Yu, 2018), and afferent heterogeneities in terms of networks receiving identical *vs*. distinct place-field inputs.

Our network simulations showed that recurrent spiking networks exhibit strong heterogeneities in firing rates and spatial information transfer, even when composed of identical units receiving identical spatial inputs. Neuronal firing rate, the extent of place-field firing, subthreshold voltage ramp, and information transfer varied markedly across all networks and were strongly influenced by trial-to-trial variability. Across all networks, consistent with prior observations in other sensory systems and in single-neuron place-cell models (Butts, 2003; Butts & Goldman, 2006; DeWeese & Meister, 1999; Montgomery & Wehr, 2010; A. Roy & Narayanan, 2021), with increasing trial-to-trial variability, the spatial location of peak information transfer transitioned from high slope regions to the peak region of the corresponding spatial tuning curve. While increasing the strength of inhibition expectedly reduced firing rates and place-field widths, peak information transfer was particularly robust across different network configuration. In networks endowed with within-type intrinsic heterogeneities, the strength of synaptic inhibition played a critical role in regulating the extent of heterogeneities in firing profiles and the differences of heterogeneous networks from their homogeneous counterparts. Strikingly, spatial information transfer in heterogeneous networks was robust to high trial-to-trial variability compared to their homogeneous counterparts. Finally, introduction of afferent heterogeneities allowed each place cell to manifest peak firing at distinct spatial locations and resulted in reduced peak information transfer and amplified diversity in spatial information transfer.

Overall, our analyses unveil degeneracy and complexity in the emergence of strong spatial tuning and efficacious spatial information transfer in recurrent networks of place cells. Among the disparate mechanisms that governed robust spatial tuning in the network were trial-to-trial variability, excitation-inhibition balance, within-type intrinsic heterogeneities, and afferent heterogeneities. Our results also highlight the critical role of neural heterogeneities in enhancing the robustness of spatial information transfer in the face of perturbations.

## METHODS

### Modelling homogeneous and heterogeneous CA3 networks with Izhikevich neuron models

The default network was constructed with 100 excitatory and 10 inhibitory neurons. To model the excitatory and the inhibitory neurons to construct the homogeneous network, we exploited the computational efficacy of abstract spiking neuronal models (Izhikevich, 2003). The spiking mechanisms in each of such neurons were governed by a system of two ordinary differential equations:

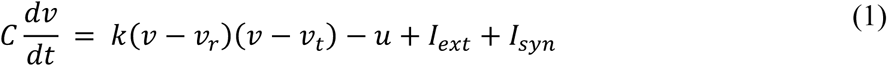

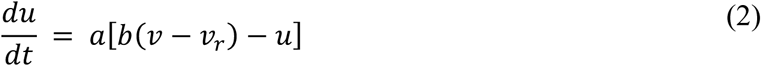

where *v* is the membrane potential, *u* is the recovery current, *C* is the membrane capacitance (set to 1), *k* defined a voltage scale factor, *v*_*r*_is the resting membrane potential, *v*_*t*_is the instantaneous threshold potential, *I*_*ext*_refers to the external current source, and *I*_*syn*_ corresponds to the total synaptic current. Parameter *a* describes the time scale of the recovery variable *u*, and *b* describes the sensitivity of the recovery variable *u* to the sub-threshold fluctuations of the membrane potential *v*. For the excitatory neurons, we adapted the tonic-bursting (TB) class of the Izhikevich neurons because bursting is one of the predominant modes of firing exhibited by the excitatory CA3 pyramidal neurons (Hemond, et al., 2008). We adapted the fast-spiking (FS) model for modelling the inhibitory neurons in the network as interneurons exhibit high-frequency firing patterns (Gloveli, et al., 2005; Golomb, et al., 2007; Kawaguchi, 2001; Markram, et al., 2004; McCormick, Connors, Lighthall, & Prince, 1985). The parameters for both the TB and the FS neurons are mentioned in Table 1. To incorporate intrinsic heterogeneity into a physiologically relevant neuronal network, the parameters *a*, *b*, *c*, and *d* of the Izhikevich model neurons were sampled randomly from a uniform distribution for each excitatory and inhibitory neuron in the network (Table 1).

**Table 1:**
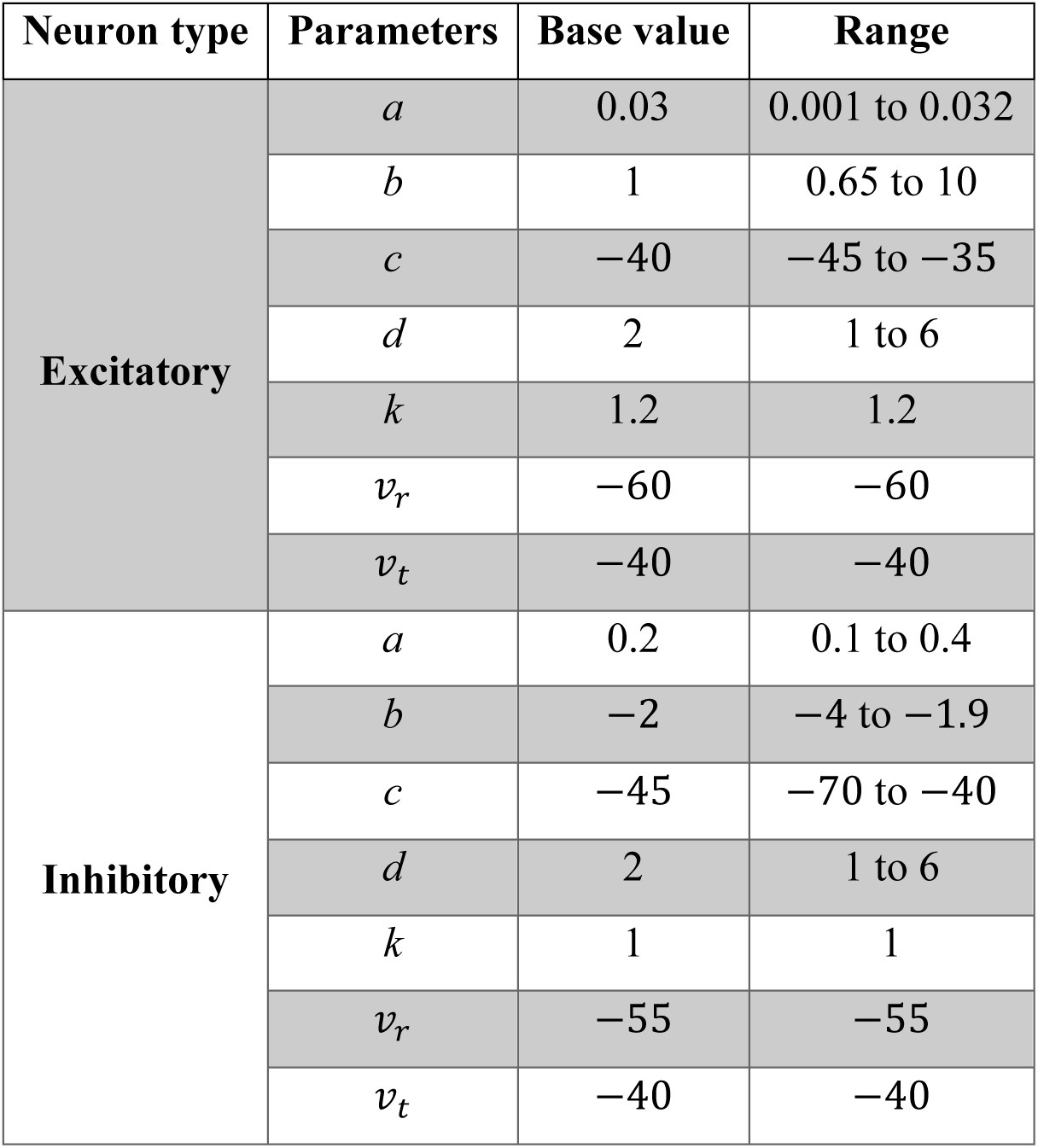
Parametric search ranges for the heterogeneous network were chosen to be around the base model value, which were adapted from (Izhikevich, 2003).

The excitatory AMPAR and the inhibitory GABAA receptor currents in each neuron in the network were modelled as Ohmic currents as:

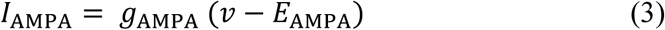

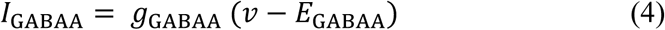

Both *g*_AMPA_ and *g*_GABAA_ followed first-order kinetics in their evolution:

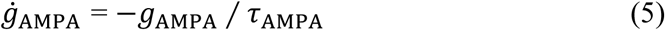

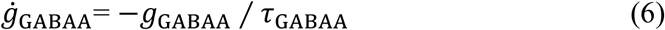

where *E*_AMPA_= 0 mV, *E*_GABAA_= –90 mV, *τ*_AMPA_= 5 ms and *τ*_GABAA_= 15 ms. Each excitatory neuron in the network received 100 spatially tuned afferent inputs. The connectivity between the neurons were specified by binary connectivity matrices, *C*_*i*j_, that were defined as per the different connection probabilities (Table 2). The connection probability for both the homogeneous and heterogeneous networks were similar (Table 2). The synaptic strengths for each of these connections for both the network types were assigned from a distinct uniform random distribution for each connection type: E → E, E → I, I → E, I → I (Table 2). Next, we introduced synaptic heterogeneities by scaling the synaptic E → I weights in the network for three sets of E–I synaptic weights in the network: 100:10, 100:20, 100:50. We introduced scaling of inhibitory weights to understand how excitation-inhibition balance contributed to the efficacy of the recurrent networks to encode information (Yu, et al., 2018).

**Table 2:** The synaptic weight distributions divided into four blocks indicating the connection probability and synaptic weight distributions for each block. ∼*U*(*x*, *y*) indicates that the numbers were derived from a uniform distribution with *x* and *y* as the lower and upper bounds, respectively. The connection probabilities reflected the CA3 network (Le Duigou, et al., 2014).

| Cell type and connections | Connection probability | Synaptic weights |
| --- | --- | --- |
| **Homogeneous Network** |  |  |
| E→E | 0.3 | $\sim U(0.1, 2)$ |
| E→I | 0.5 | $\sim U(0.01, 0.09)$ |
| I→E | 0.5 | $\sim U(-1, -0.01)$ |
| I→I | 0.7 | $\sim U(-0.1, -0.01)$ |
| **Heterogeneous Network** |  |  |
| E→E | 0.3 | $\sim U(0.1, 1)$ |
| E→I | 0.5 | $\sim U(0.01, 0.09)$ |
| I→E | 0.5 | $\sim U(-0.02, -0.01)$ |
| I→I | 0.7 | $\sim U(-0.1, -0.01)$ |

### Singular place field like input to the network for afferent synaptic stimulation

To investigate the relationship between spatial tuning and spatial information transfer in the network, we stimulated all the excitatory neurons with place field-like inputs through randomly distributed 100 afferent synapses. The synaptic weights for each afferent input to each excitatory neuron was chosen randomly from a uniform distribution (∼*U*(1,3)). The Gaussian modulated co-sinusoidal function was fed as an probabilistic afferent input (Basak & Narayanan, 2018, 2020; A. Roy & Narayanan, 2021; Seenivasan & Narayanan, 2020) to all of the 100 synapses across each of the excitatory neuron to mimic the place field like inputs to these neurons. The frequency of the Gaussian input was set to 8 Hz, representative of the theta oscillations accompanying the place cell response when the animal traverses any spatial location. The probability of the occurrence of the input to each of the synapses at any given time point was driven by:

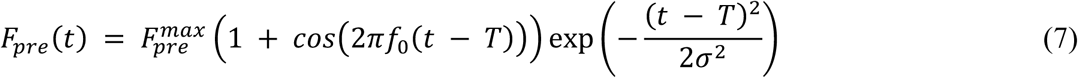

where *T* corresponds to the place field center, 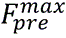 is the maximal input firing rate, *f*_0_ is the frequency of the cosine (8 Hz), and *σ* corresponds to the standard deviation of the Gaussian function that controls the extent of the place field input (0.5 s). The resultant synaptic current produced post-synaptic depolarization and action potentials that were reflective of place field activity profiles. In our simulations, we assumed that the virtual animal was traversing at a constant velocity, implying the equivalence of space and time variables (*t* in eqn. (7)). The afferent synaptic stimulation was fixed for all the different sets of synaptic E–I ratios mentioned above – 100:10, 100:20 and 100:50 respectively.

### Distinct place field-like afferent synaptic inputs to the network

The single-place field-like input does not take into account the different presynaptic neurons, each endowed with heterogeneous place field locations and differential synaptic weights impinging onto the postsynaptic neuron (Bittner, et al., 2015; Bittner, Milstein, Grienberger, Romani, & Magee, 2017; Dong, Madar, & Sheffield, 2021; Grienberger, Milstein, Bittner, Romani, & Magee, 2017). We considered the scenario where each excitatory neuron received inputs not as a single spatially tuned input, but from a multitude of neurons with each endowed with different spatial firing profiles. In implementing this, we subjected each of the excitatory neurons in the network to spatially modulated inputs from 10 different presynaptic neurons such that each of them had spatial tuning profiles at distinct locations (A. Roy & Narayanan, 2021).

### Quantification of instantaneous firing rate during place field traversal

The spike times were derived from the voltage profile based on when the voltage crossed the set threshold of –20 mV on the rising phase of the voltage response. The derived spike times were then discretized into binary time series consisting of 0 and 1, where 0 implied the absence of spike and 1 implied the presence of spike in the temporal window (bin width = 1 ms). The spike train was then convolved with a Gaussian kernel (*σ* = 200 ms) to obtain the smooth instantaneous firing rate. A set of measurements (Basak & Narayanan, 2018, 2020; A. Roy & Narayanan, 2021; Seenivasan & Narayanan, 2020) were quantified from the instantaneous firing rate profile for all the excitatory neurons across three sets of synaptic E:I ratios – (i) the maximum firing rate of the place cell (*F*_*max*_), (ii) full-width at half maximum (*FWHM*) defined as the temporal distance between the two-maximal half points (on either side of the center) of the curve. These measures are significant indicators of spatial tuning in place cell responses, where a high *F*_*max*_and lower *FWHM* would translate to a sharply tuned place cell response (Basak & Narayanan, 2018, 2020; A. Roy & Narayanan, 2021). As the animal traverses along any place field, the place cells have been reported to generate characteristic sub-threshold ramp voltages which is reflected in their intracellular voltage profiles (Bittner, et al., 2017; Harvey, et al., 2009). To evaluate the presence of such distinct ramp voltages in the place cells in the network, we subjected the place cell voltage to a wide median filter of 0.5 s (Basak & Narayanan, 2018, 2020; A. Roy & Narayanan, 2021; Seenivasan & Narayanan, 2020) such that this eliminated the spikes and revealed the presence or absence of a voltage ramp. Following this, the ramp voltage amplitude was computed as the maximum value of the voltage attained by the ramp.

### Introduction of trial-to-trial variability in the place cell responses in the networks

To achieve variability in the responses of the place cells in the networks, different levels of additive noise were introduced to the pre-synaptic firing rate profile (eqn. (7)) for each of the synapses. The noise was introduced in the form of an additive Gaussian white noise (AGWN) denoted by:

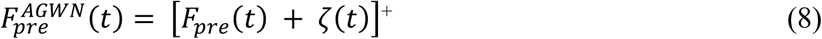

[*F*]^+^ = max(*F*, 0) corresponds to the rectification to avoid negative firing rates, *ζ*(*t*) corresponds to AGWN with mean zero and standard deviation *σ*_*noise*_. Different values of *σ*_*noise*_with *F*_*pre*_(*t*) were representative of the trial-to-trial variability which was utilized to assess the impact of the trial-to-trial variability in symmetric place-field firing profiles. Since the rectification governed the overall firing rate and not the noise term, this type of formulation allowed for both negative and positive modulation of *F*_*pre*_(*t*). The AGWN introduced trial-to-trial variability across all the locations in the spatial field, irrespective of the strength of the afferent synaptic activity, and thus translates to activity-independent variability (A. Roy & Narayanan, 2021).

### Spatial information transfer within a place field by the CA3 network: Stimulus-specific information metrics

To quantify spatial information transfer across multiple spatial locations, we computed stimulus-specific information (SSI) for all the spatially tuned neurons in the networks. Spatial information transfer was derived from the SSI computed across 25 independent trials of network simulations spanning the entire traversal of the linear spatial arena. To calculate the SSI value, the firing rate response and the stimulus were divided into 40 and 80 bins respectively (Butts, 2003; Butts & Goldman, 2006; Montgomery & Wehr, 2010; A. Roy & Narayanan, 2021). The specific information of a response is given by:

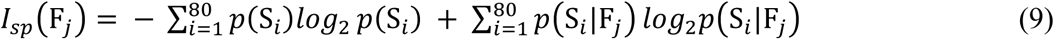

where *p*(F_j_) denoted the probability of the firing rate in the *j^th^* response bin and *p*(S_*i*_|F_j_) corresponded to the conditional probability for the stimulus in the *i^th^*bin given that the firing rate was in the *j*^th^ response bin. The first term in (eqn. (9)) corresponds to the stimulus entropy and the second term in (eqn. (9)) corresponds to the total noise entropy. The stimulus-specific information was then computed as:

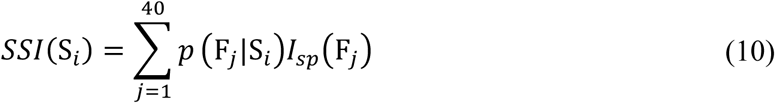

where *p*(F_j_|S_*i*_) defined the conditional probability of the firing rate in the *j*^th^ response bin given the *i*^th^ stimulus was presented. Therefore, specific information defines the reduction in the uncertainty about the spatial location gained by a particular firing rate response CF_j_H. SSI corresponds to the average specific information of the responses that manifest when a particular spatial stimulus is presented. The bias in the *I*_*sp*_was corrected using the Treves-Panzeri correction method (Montgomery & Wehr, 2010; Panzeri, Senatore, Montemurro, & Petersen, 2007; Treves & Panzeri, 1995):

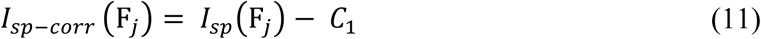

where 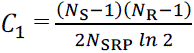, *N*_R_ is the number of response bins, *N*_S_ is the total number of stimulus bins, and N_SRP_ represents the total number of stimulus-response pairs. This was done to compensate for the biases that are associated with the information measures due to the finite number of trials in the experiment, which further leads to the under sampling of the relevant probability distributions. We computed the SSI metric for each of the 100 excitatory neurons across their responses to the place field-like inputs. To quantify the information transfer across different sets of trial-to-trial variability and for different regimes of synaptic E:I balance, we computed two quantities: (i) *SSI peak*, which corresponds to the maximal value at the center of the place field, (ii) *SSI FWHM*, which corresponds to the full-width at half maxima of the SSI tuning curve (A. Roy & Narayanan, 2021).

### Computational details

All simulations were performed using the NEURON programming environment (Carnevale and Hines, 2006), at 34°C. The simulation step size was set as 25μs. Data analysis and plotting of graphs were done by using custom written codes in MATLAB and IGOR Pro environment (WaveMetrics Inc., Portland, OR, USA). Statistical analysis was performed in R (www.R-project.org). All the data points across all the stated simulations have been reported and represented in the form of bee swarm and scatter plots (wherever deemed necessary) in order to avoid any misleading interpretations from just reporting the overall summary statistics (Marder & Taylor, 2011; Mittal & Narayanan, 2018; Rathour & Narayanan, 2014; A. Roy & Narayanan, 2021).

## RESULTS

The question that we addressed in this study is how spatial information transfer occurs in homogeneous *vs*. heterogeneous CA3 networks of spatially tuned neurons embedded within a recurrent network. In addressing this question, we systematically assessed the impact of three different forms of neural-circuit heterogeneities (Dahmen, et al., 2025; Mishra & Narayanan, 2019, 2021b; Mittal & Narayanan, 2021; Saini & Narayanan, 2025; Santhosh & Narayanan, 2025) and trial-to-trial variability (Butts, 2003; Butts & Goldman, 2006; A. Roy & Narayanan, 2021) on firing rate tuning profiles and information transfer:

1. **Trial-to-trial variability**: Different levels of trial-to-trial variability introduced through additive Gaussian white noise;
2. **Synaptic heterogeneities**: Afferent synaptic heterogeneities in terms of synaptic weights of afferent inputs onto individual neurons and local synaptic heterogeneities in terms of different levels of excitation:inhibition (E:I) balance in the local network;
3. **Intrinsic heterogeneities**: Defined by networks constructed of neurons with identical *vs*. heterogeneous intrinsic properties; and
4. **Afferent heterogeneities**: Defined by networks receiving identically tuned *vs*. distinctly tuned afferent inputs.

### Network configuration, E:I balance, trial-to-trial variability, and information metrics

We first built a homogeneous network comprising of 100 excitatory and 10 inhibitory neurons with connection probabilities reflecting the CA3 network (Fig. 1*A–B*; Tables 1–2). Excitatory and inhibitory neurons modelled as tonic bursting and fast spiking neurons, respectively (Fig. 1*A*). In this homogeneous network, all excitatory neurons were identical to one another, as were the inhibitory neurons, with neuronal parameters set to their respective base values (Table 1). We stimulated the network through 100 afferent synapses to each excitatory cell with place field-like inputs to generate spatially tuned responses for each of the place cells (Fig. 1*B*). All afferent inputs were identically tuned to space. We repeated this procedure for 25 independent trials and for three different noise levels (low, medium, and high) through an additive Gaussian white noise (AGWN) incorporated into the frequency of synaptic stimulation. The excitation-inhibition (E:I) balance in the network was controlled by altering inhibitory synaptic weights with all simulations repeated for three distinct values of E:I ratio, namely 100:10, 100:20, and 100:50. Spatial information transfer for each excitatory neuron was computed using stimulus specific information (SSI) metrics across the three different noise levels and three different values of E:I ratio, constituting a total of 22,500 (100 × 25 × 3 × 3) network simulations for every network configuration.

**Figure 1:**
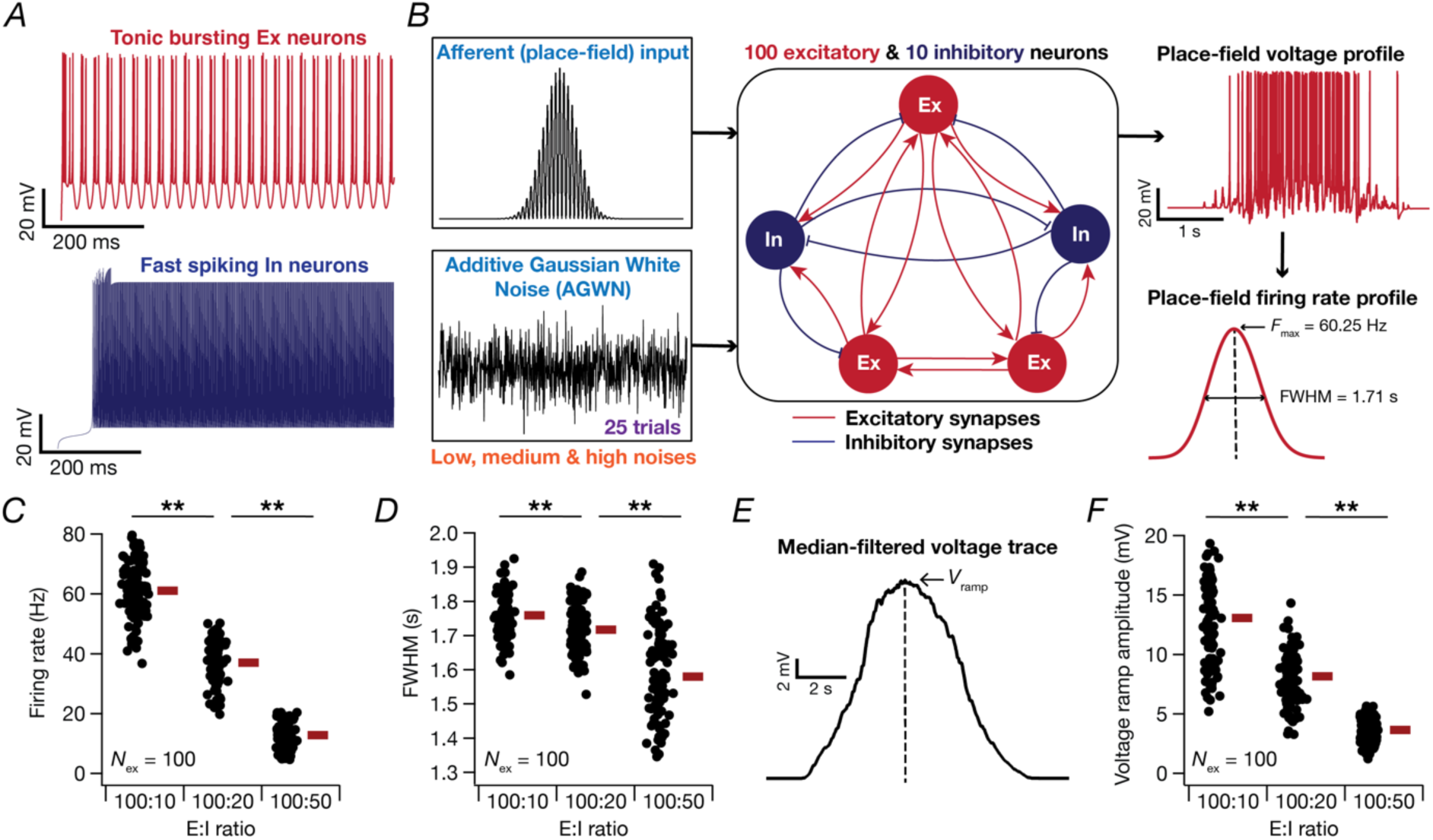
Synaptic excitation-inhibition balance regulates place cell characteristics of principal neurons in the homogeneous CA3 network. (*A*) *Top*, Voltage trace of the excitatory neuron (modelled as tonic bursting neuron) in response to 500 pA current for 1 s and *bottom*, Voltage trace of the inhibitory neuron (modelled as fast spiking neuron) in response to 43 pA current for 1 s. (*B*) Depiction of the CA3 network with recurrent connectivity between the neurons (Table 2 shows connection probabilities). Red: excitatory connections and Blue: inhibitory connections. Left panels depict the inputs to the network. *Top,* The afferent input is specified as a Gaussian modulated sinusoid defining the probability distribution of activating 100 independent presynaptic inputs arriving onto each of the excitatory neurons in the recurrent network during a place field traversal. *Bottom,* To introduce trial-to-trial variability in the responses across each of the neurons in the network, the place field-like inputs were introduced along with an additive Gaussian white noise (AGWN) for 3 different levels of noise (low, medium, and high) by varying the standard deviation *σ*_*noise*_. Right panels depict the responses of neurons in the network. *Top*, Example voltage trace of the excitatory neuron in response to the afferent synaptic activation. *Bottom*, Instantaneous firing rate curve derived from the voltage response. All excitatory neurons in the network were stimulated with place field-like inputs along with the AGWN across 25 independent trials for each of the 3 different noise levels and different E–I levels. (*C–D*) Impact of different synaptic excitation-inhibition weight ratios on the peak firing frequency, *F*_*max*_ (*C*) and full-width at half maximum, *FWHM* (*D*) of the 100 excitatory neurons in the network. The red bars represent the respective median values for each of the scenarios, *F*_*max*_: Wilcoxon signed rank test, 100:10 *vs*. 100:20 *p* = 2.2 × 10^−16^, 100:20 *vs*. 100:50 *p* = 2.2 × 10^−16^, 100:10 *vs*. 100:50 *p* = 2.2 × 10^−16^, *FWHM*: Wilcoxon signed rank test, 100:10 *vs*. 100:20 *p* = 4.04 × 10^−5^, 100:20 *vs*. 100:50 *p* = 3.95 × 10^−13^, 100:10 *vs*. 100:50 *p* = 2.2 × 10^−16^ (*E*) Voltage profile in (*B*) was filtered using a median filter of 0.5 s window to obtain the sub-threshold voltage ramp, which is a characteristic hallmark of the intracellular voltage during a place field traversal. *V*_*ramp*_represents the peak of the voltage ramp. (*F*) Impact of different synaptic excitation-inhibition weight ratios on the ramp voltage amplitude of the 100 excitatory neurons in the network. Kruskal Wallis test, *p* = 0.481, Wilcoxon signed rank test, 100:10 *vs*. 100:20 *p* = 2.2 × 10^−16^, 100:20 *vs*. 100:50 *p* = 2.2 × 10^−16^, 100:10 *vs*. 100:50 *p* = 2.2 × 10^−16^.

### Heterogeneities in place-field characteristics of a homogeneous CA3 network receiving identical afferent inputs

The firing rate profile of a place cell as a function of space generates a spatial tuning curve (Fig. 1*B*). The spatial tuning curve has been extensively studied for single place cells using morphologically realistic models endowed with different biophysical heterogeneities (A. Roy & Narayanan, 2021). To study the spatial tuning features of place cells embedded in a recurrent network, we first incorporated 100 afferent synapses randomly on each of the 100 excitatory neurons in the homogeneous CA3 network. The synaptic weights for each of the afferent synapses in an excitatory neuron chosen randomly from a uniform distribution, ensuring that afferent synaptic weights were heterogeneous. Afferent synapses were activated in a probabilistic manner with a Gaussian modulated co-sinusoidal function, with the field center being at *T* = 5 s. We assumed that the velocity of the virtual animal traversing a linear arena was constant, thus providing equivalence of space and time taken for the traversal.

We obtained the voltage responses corresponding to these stimulus conditions and computed the instantaneous firing rate profiles (Fig. 1*B*) for all the 100 stimulated excitatory neurons in the network. A sharply-tuned place field response appears as a smooth Gaussian function, with maximal firing at the place field center accompanied by sharply reducing responses on either side of the center (Fig. 1*B*). We repeated these analyses for the three different sets of synaptic E:I strengths and quantified place-cell responses using peak firing rate, *F*_*max*_ and full width at half-maxima, *FWHM* for each neuron in the network (Fig. 1*C–D*). As subthreshold ramp voltages are representative of place-cell firing (Harvey, et al., 2009; Li, Briguglio, Romani, & Magee, 2024), we computed subthreshold voltage ramps (Fig. 1*E*) by median filtering the voltage response (Basak & Narayanan, 2018, 2020; A. Roy & Narayanan, 2021). We calculated the peak ramp voltage *V*_*ramp*_ from the subthreshold voltage ramps for all the place-cell responses (Fig. 1*F*). With increasing inhibition, we found an expected progressive reduction in the firing rate (Fig. 1*C*), extent of place-field firing (Fig. 1*D*), and the subthreshold voltage ramp (Fig. 1*F*). Importantly we noted pronounced heterogeneities across all three quantities (Fig. 1*C–D*, Fig. 1*F*). However, the maximal firing for all neurons occurred for the place field center (*T* = 5 s) because of the homogeneous afferent inputs that all neurons received in this set of simulations.

Together, these observations unveiled heterogeneities in the firing rate, extent of place-field, and subthreshold voltage ramp amongst neurons within the same homogeneous network. We also demonstrate strong dependence of these quantities on synaptic inhibition, with enhanced inhibition resulting in reductions in firing rate, voltage ramp amplitude, and the extent of place-field firing.

### Trial-to-trial variability in place-cell responses regulates spatial tuning curve and spatial information transfer in homogeneous CA3 networks receiving identical afferent inputs

The spatial tuning curve of place cells serves as a substrate to encode information about the spatial locations that an animal traverses. The place field center elicits the peak firing response, and the responses reduce significantly for all the corresponding locations on either side of the peak (Fig. 1*B*). An important question associated with the spatial tuning curve is on how much information about each location is conveyed by the firing rates. Does the maximal spatial information transfer occur at the peak of the spatial tuning curve or at the high-slope regions? Previous studies have reported information transfer to be strongly dependent on trial-to-trial variability across different sensory modalities (Butts & Goldman, 2006; Montgomery & Wehr, 2010; A. Roy & Narayanan, 2021). To understand the impact of trial-to-trial variability on the spatial tuning curves in a recurrent neuronal network, we introduced an additive Gaussian noise term to the afferent input rate. We introduced three different levels of additive Gaussian noises (low, medium, and high) governed by the variance term in the Gaussian. We obtained the spatial tuning profiles of place cells in the network for the 3 different levels of noise and for 3 levels of E:I strengths (Fig. 2).

**Figure 2:**
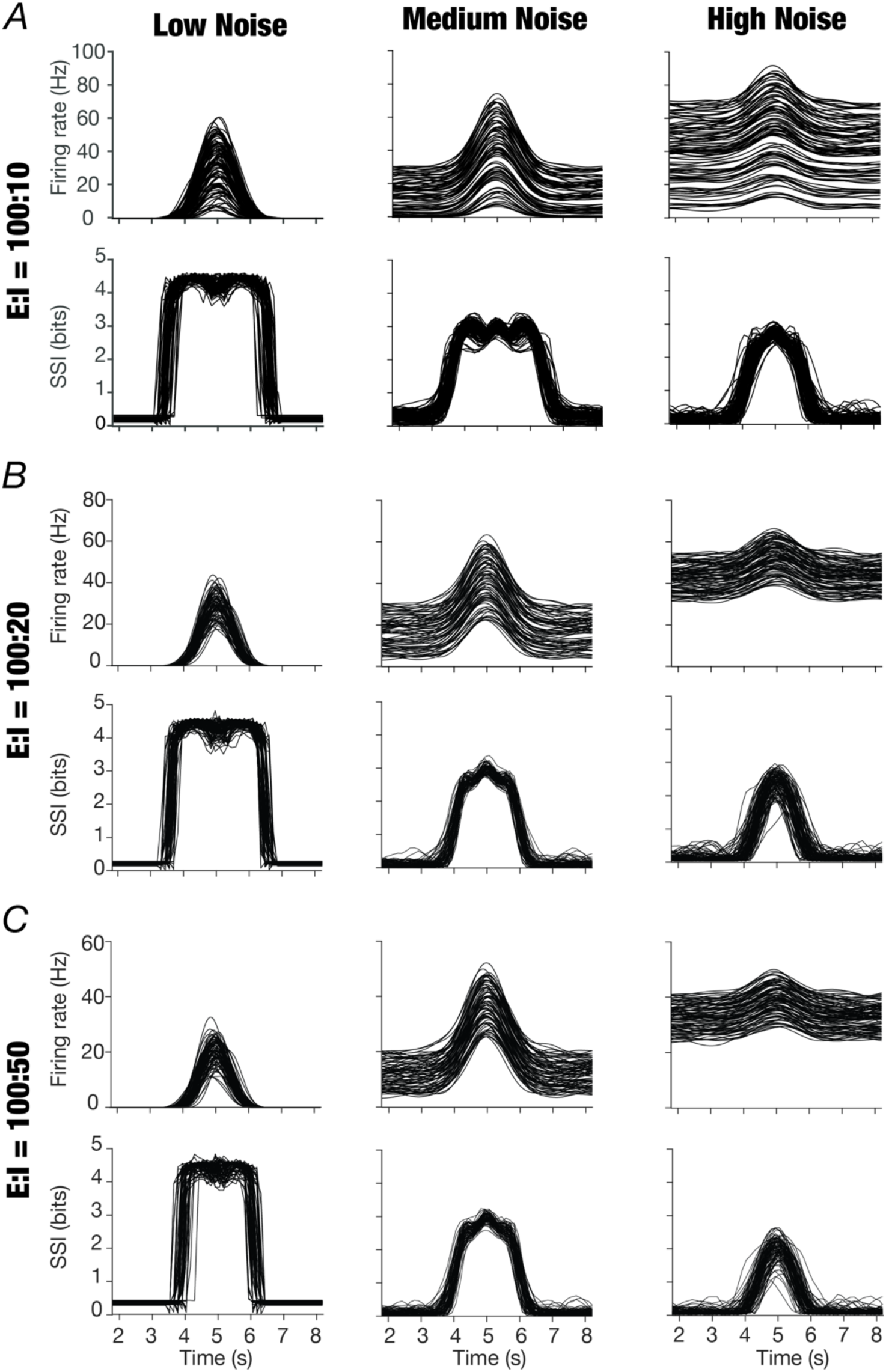
Impact of trial-to-trial variability and excitation-inhibition balance on spatial information transfer by the place cells in homogeneous CA3 networks. (*A–C*) Mean firing rate (averaged across all 25 trials) and the corresponding stimulus-specific information profiles for the 100 excitatory neurons in the network for excitation-inhibition (E:I) ratio of 100:10 (*A*), 100:20 (*B*), and 100:50 (*C*). The columns represent the low, medium and high trial-to-trial variability introduced by AGWN. *σ*_*noise*_ values for *Low*: 1 × 10^−6^ Hz^2^, *Medium*: 1 × 10^−2^ Hz^2^, *High*: 1 Hz^2^.

We observed an expected reduction in the average firing rate of neurons in the network with increasing synaptic inhibition (Fig. 2). For low levels of trial-to-trial variability, the out-of-field firing was low and the tuning curve showed a sharp Gaussian profile across all levels of inhibition. Under high trial-to-trial variability, both the in-field (within the place field) and the out-of-field peak firing rates increased significantly resulting in a relative loss of spatial tuning, with amplified peak firing rates as compared to that of the low and medium levels of trial-to-trial variability (Fig. 2). Importantly, irrespective of the level of excitation-inhibition balance, increasing trial-to-trial variability (through enhanced variance of the AGWN) resulted in notable reductions in spatial information transfer. This may be observed from the reduction in the peak value of the SSI profile with increasing noise (Fig. 2). In addition to this reduction in spatial information transfer with increasing trial-to-trial variability, we also noted that the relationships between the firing rate tuning curves and SSI metrics (across different levels of trial-to-trial variability) were similar to earlier observations from different sensory modalities and single-neuron place-field analyses (Butts & Goldman, 2006; Montgomery & Wehr, 2010; A. Roy & Narayanan, 2021). Specifically, we observed that the SSI profile showed peaks at the maximum-slope regions of the firing-rate tuning curve when trial-to-trial variability was low (Fig. 2; low noise). With increase in trial-to-trial variability, the peak locations on the SSI profile progressively shifted to the peak location of the tuning curve (Fig. 2; high noise).

Together, these observations demonstrate a strong dependence of firing rate profiles and spatial information transfer of neurons in a homogeneous network on trial-to-trial variability, consistent with earlier observations with detailed single neuron models of place cells in the CA1 region of the hippocampus (A. Roy & Narayanan, 2021).

### Heterogeneities in the regulation of spatial information transfer by trial-to-trial variability in place-cell responses in the homogeneous recurrent network receiving identical afferent inputs

Although all the excitatory neurons in the network are intrinsically identical, we found the SSI tuning curves to be heterogeneous across different neurons for all E:I strengths, reflecting the contribution of the different synaptic weights in the homogeneous recurrent network (Fig. 2). To systematically assess this heterogeneity, we calculated two independent SSI measures: *SSI peak*, which defined the information transferred at the center of the place fields (*T*=5 s) and *SSI FWHM*, which quantified the width of the SSI tuning curve (A. Roy & Narayanan, 2021). We derived these quantifications from the SSI tuning profiles for each of the 100 excitatory neurons, computed from 25 trials for each of the 3 noise levels and 3 synaptic E:I strengths (Fig. 3). We observed pronounced heterogeneities in spatial information transfer across neurons in the homogeneous network, for different synaptic E:I strengths and for different levels of trial-to-trial variability (Fig. 3). While *SSI peak* changed negligibly with increased inhibition, *SSI FWHM* reduced significantly with increase in the strength of synaptic inhibition (Fig. 3). In contrast, across all values for E:I balance, both *SSI peak* and *SSI FWHM* significantly and progressively reduced with gradual increase in trial-to-trial variability (Fig. 3).

**Figure 3:**
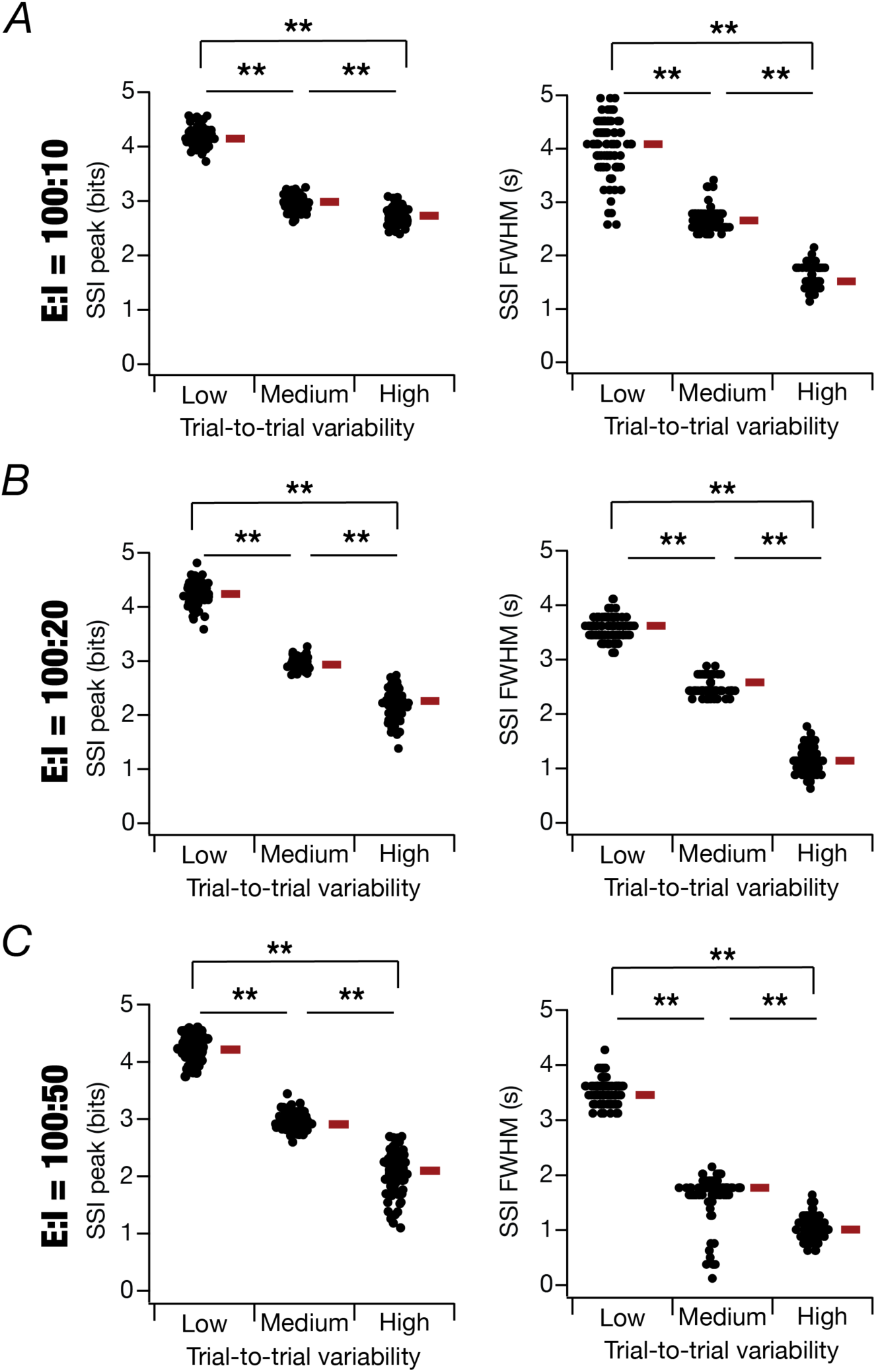
Spatial information transfer reduces with increase in trial-to-trial variability across all levels of excitation-inhibition balance in homogeneous CA3 networks. (*A*) Stimulus-specific information (SSI) value at the center of the place field (*T*=5 s), *SSI peak* and the full width at half-maxima, *SSI FWHM* computed for E:I synaptic strength ratio = 100:10. *SSI peak*: Kruskal Wallis test, *p* = 0.481, Wilcoxon Rank Sum test, Low *vs*. Medium *p* = 2.2 × 10^−16^, Medium *vs*. High *p* = 2.2 × 10^−16^, Low *vs*. High *p* = 2.2 × 10^−16^, *SSI FWHM*: Kruskal Wallis test, *p* = 0.48, Wilcoxon Rank Sum test, Low *vs*. Medium *p* = 4.9 × 10^−14^, Medium *vs*. High *p* = 2.2 × 10^−16^, Low *vs*. High *p* = 2.2 × 10^−16^. (*B*) *SSI peak* and *SSI FWHM* computed for E:I synaptic strength ratio = 100:20. *SSI peak*: Kruskal Wallis test, *p* = 0.48, Wilcoxon Rank Sum test, Low *vs*. Medium *p* = 2.2 × 10^−16^, Medium *vs*. High *p* = 2.2 × 10^−16^, Low *vs*. High *p* = 2.2 × 10^−16^, *SSI FWHM*: Kruskal Wallis test, *p* = 2.2 × 10^−16^, Wilcoxon Rank Sum test, Low *vs*. Medium *p* = 2.2 × 10^−16^, Medium *vs*. High *p* = 2.2 × 10^−16^, Low *vs*. High *p* = 2.2 × 10^−16^. (*C*) *SSI peak* and *SSI FWHM* computed for E:I synaptic strength ratio = 100:50, *SSI peak*: Kruskal Wallis test, *p* = 0.35, Wilcoxon Rank Sum test, Low *vs*. Medium *p* = 2.2 × 10^−16^, Medium *vs*. High *p* = 2.37 × 10^−16^, Low *vs*. High *p* = 2.2 × 10^−16^, *SSI FWHM*: Kruskal Wallis test, *p* = 0.37, Wilcoxon Rank Sum test, Low *vs*. Medium *p* = 2.2 × 10^−16^, Medium *vs*. High *p* = 2.2 × 10^−16^, Low *vs*. High *p* = 2.2 × 10^−16^. *σ*_*noise*_ values for *Low*: 1 × 10^−6^ Hz^2^, *Medium*: 1 × 10^−2^ Hz^2^, *High*: 1 Hz^2^.

Together, these observations emphasized the manifestation of heterogeneities in spatial information transfer even in a homogeneous network, with strong dependence of the information transfer on trial-to-trial variability. Although the width of the SSI profile reduced with increase in inhibition owing to a reduction in the width of the firing rate profile (Fig. 2), there was no major impact of inhibitory balance on the peak value of spatial information transferred.

### Heterogeneities in the place cell responses and characteristic place field properties of the heterogeneous network receiving identical afferent inputs under different regimes of synaptic E:I balance

Thus far, our analyses have been restricted to a homogeneous CA3 network, in which all excitatory neurons were identical to one another, as were the inhibitory neurons. To understand the impact of widespread heterogeneities in neuronal intrinsic properties on spatial tuning and information transfer, we introduced heterogeneities into the excitatory and inhibitory neuronal population. Specifically, instead of setting neuronal parameters of all excitatory and inhibitory neurons in the network to their respective base values (Table 1), we randomly picked unique parametric values for each neuron in the network. Excitatory and inhibitory neurons had different distributions from where their unique parametric values were randomly picked from (Table 1). The choice of different parameters resulted in a heterogeneous population of excitatory (Fig. 4*A*) and inhibitory (Fig. 4*B*) neurons. While the heterogeneous network was constructed with neurons with non-repeating parameters, all other network parameters including the number of neurons, their local connectivity, and afferent synaptic connectivity (Fig. 4*C*) were identical to those of the homogeneous network. We then simulated this heterogeneous recurrent network in a manner identical to the homogeneous network. We characterized place cell responses by computing the instantaneous firing rates for all the 100 excitatory neurons and computed *F*_*max*_, *FWHM*, and *V*_*ramp*_respectively. We repeated this procedure for heterogeneous networks endowed with different E:I balance.

**Figure 4:**
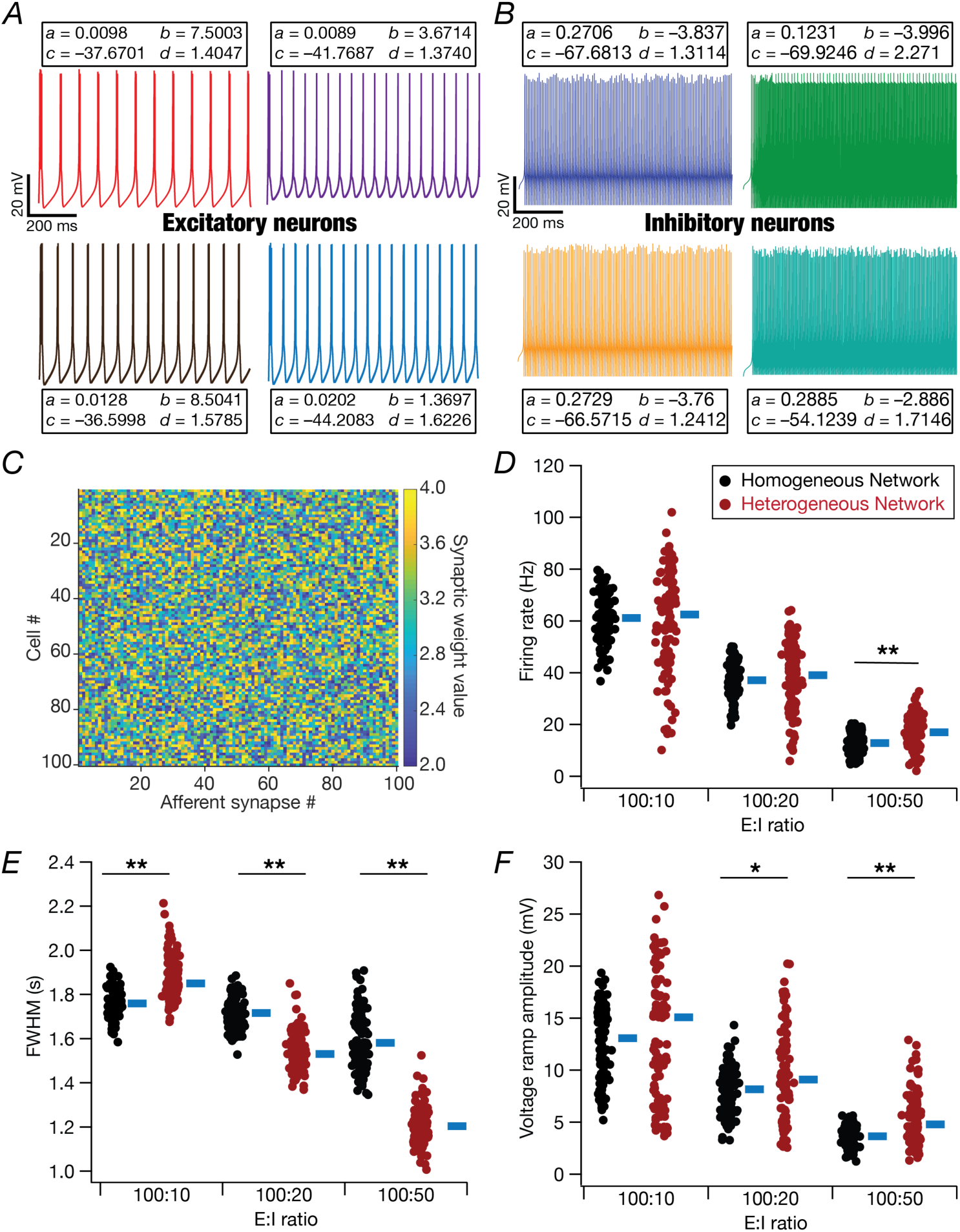
Comparison of place-field characteristics between homogeneous and heterogeneous network neurons. (*A*–*B*) Voltage responses of 4 representative (neuron #12, 47, 66, 90) excitatory neurons (A) and 4 representative (neuron #2, 4, 6, 10) inhibitory neurons (B) to pulse current injections, showing heterogeneity of intrinsic properties of these neurons in a heterogeneous network. The inset boxes associated with each trace refers to the parameters for each of the neurons which were picked from a random uniform distribution. (*C*) Distribution of the afferent synaptic weights for 100 afferent synapses impinging onto each of the excitatory neurons in the heterogeneous network. (*D*–*F*) Impact of different synaptic E:I strengths on the peak firing frequency, *F*_*max*_(*D*), full-width at half maximum, *FWHM* of the firing rate profile (*E*) and sub-threshold ramp voltage, *V*_*ramp*_ (*F*) of the 100 excitatory neurons in the homogeneous and heterogeneous networks. The blue bars represent the respective median values for each of the measures. Statistical comparisons for heterogeneous networks, *F*_*max*_: Kruskal Wallis test, *p* = 2.2 × 10^−16^, Wilcoxon rank sum test, 100:10 *vs*. 100:20 *p* = 8.7 × 10^−15^, 100:20 *vs*. 100:50 *p* = 2.2 × 10^− 16^, 100:10 *vs*. 100:50 *p* = 2.2 × 10^−16^, *FWHM*: Kruskal Wallis test, *p* = 2.2 × 10^−16^, Wilcoxon rank sum test, 100:10 *vs*. 100:20 *p* = 2.2 × 10^−16^, 100:20 *vs*. 100:50 *p* = 2.2 × 10^−16^, 100:10 *vs*. 100:50 *p* = 2.2 × 10^−16^, *V*_*ramp*_: Kruskal Wallis test, *p* = 2.2 × 10^−16^, Wilcoxon Rank Sum test, 100:10 *vs*. 100:20 *p* = 9.1 × 10^−6^, 100:20 *vs*. 100:50 *p* = 2.1 × 10^−12^, 100:10 *vs*. 100:50 *p* = 2.2 × 10^−16^. Statistical comparisons between homogeneous *vs.* heterogeneous networks: *F*_*max*_: Kruskal Wallis test, *p* = 2.2 × 10^−16^; Wilcoxon rank sum test, 100:10, *p* = 0.54; 100:20, *p* = 0.08; 100:50, *p* = 2.6 × 10^−7^. *FWHM*: Kruskal Wallis test, *p* = 2.2 × 10^−16^; Wilcoxon Rank Sum test, 100:10, *p* = 2.2 × 10^−16^; 100:20, *p* = 2.2 × 10^−16^; 100:50, *p* = 2.2 × 10^−16^. *V*_*ramp*_: Kruskal Wallis test, *p* = 2.2 × 10^−16^; Wilcoxon rank sum test, 100:10, *p* = 0.37; 100:50, *p* = 0.02; 100:10 *vs*. 100:50 *p* = 6.1 ×10^−7^.

Similar to our results with homogeneous networks, we found a reduction in each of *F*_*max*_, *FWHM*, and *V*_*ramp*_with increasing level of synaptic inhibition in the heterogeneous recurrent network (Fig. 4*D–F*). Importantly, we consistently observed an increase in the ranges of each of *F*_*max*_, *FWHM*, and *V*_*ramp*_ in heterogeneous networks compared to their homogeneous counterparts (Fig. 4*D–F*). The median values of firing rates were comparable between the homogeneous and heterogeneous networks across all values for E:I balance, although we observed a small yet significant increase in the firing rate of heterogeneous networks when E:I balance was set at 100:50 (Fig. 4*D*). Whereas *FWHM* was higher in heterogeneous networks with lower inhibition, with increased inhibition, the heterogeneous networks showed lower *FWHM* of their firing profile compared to homogeneous networks (Fig. 4*E*). Finally, the median values of voltage ramp amplitude were consistently higher in heterogeneous networks than in their homogeneous counterparts across all values for E:I balance, with significant differences observed for E:I balance values of 100:20 and 100:50 (Fig. 4*F*).

Together, these observations showed that neurons with heterogeneous intrinsic properties manifested heterogeneities in their place-field firing profiles when placed within a recurrent network. Importantly, the extent of heterogeneities in firing profiles and the differences of heterogeneous networks from their homogeneous counterparts were critically regulated by the strength of synaptic inhibition.

### Heterogeneous networks receiving identical afferent inputs manifested enhanced spatial information transfer compared to their homogeneous counterparts with similar levels of trial-to-trial variability

We explored the impact of trial-to-trial variability in the spatial tuning profiles of the intrinsically distinct place cells in the heterogeneous network. We computed the spatial tuning curves across 25 independent trials by introducing AGWN to the pre-synaptic activity by varying the *σ*_*noise*_ for three different noise levels and for the 3 different regimes of synaptic E:I balance. Similar to our earlier observations for the homogeneous networks with increasing trial-to-trial variability (Fig. 2), we found an increase in off-field firing of individual neurons in the heterogeneous network for all levels of E:I balance (Fig. 5). However, unlike the homogeneous network where there was a shift in the peak firing rate with increasing trial-to-trial variability, we found the ranges of the peak firing rates to be comparable across different degrees of trial-to-trial variability (Fig. 5). Increasing synaptic inhibition resulted in reductions in overall firing rates and in the width of the tuning curves, but the ranges of peak firing rates across different degrees of trial-to-trial variability were comparable for a given level of E:I balance (Fig. 5).

**Figure 5.**
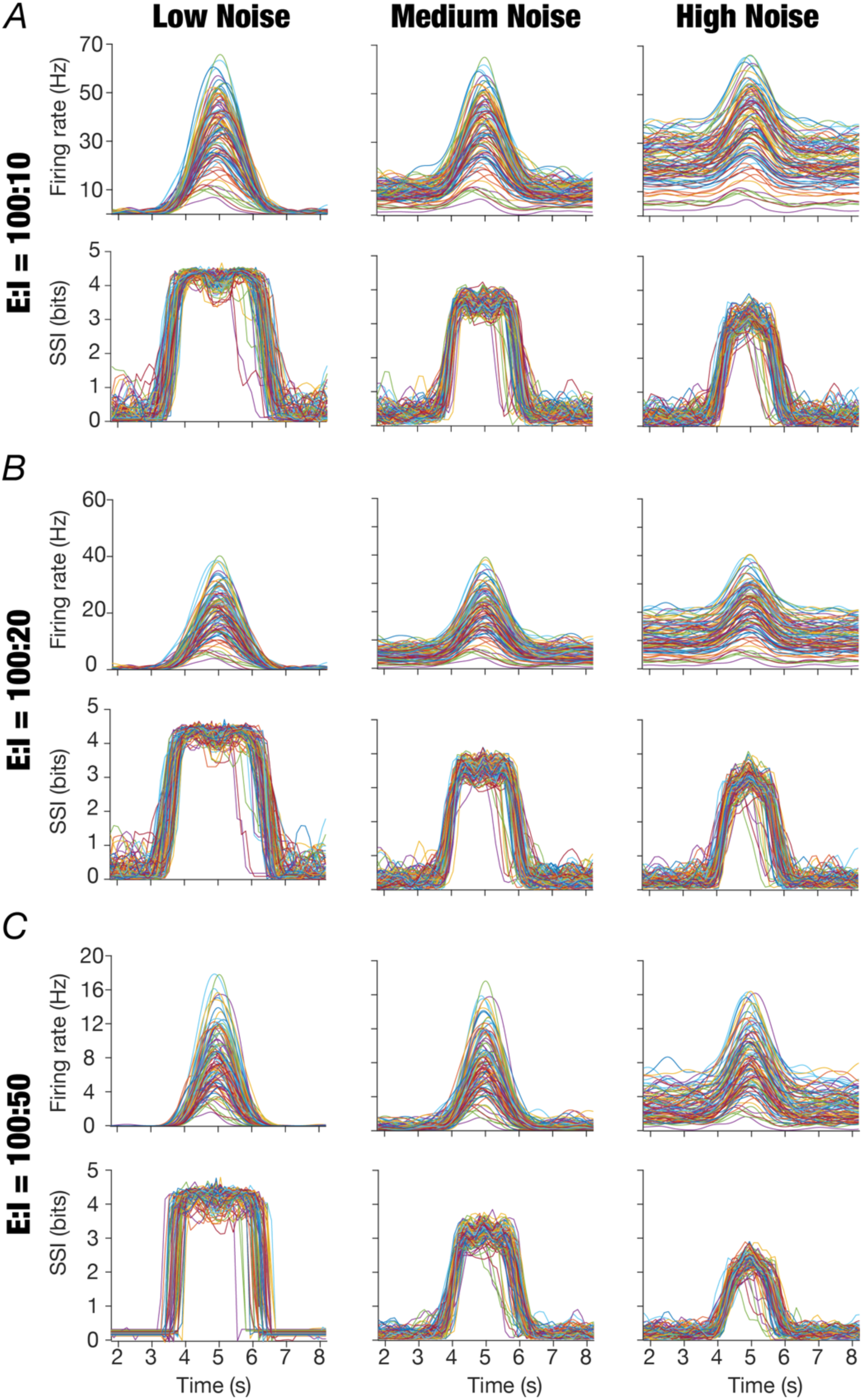
Impact of trial-to-trial variability and excitation-inhibition balance on spatial information transfer by the place cells in heterogeneous CA3 networks. (*A–C*) Mean firing rate (averaged across all 25 trials) and the corresponding stimulus-specific information profiles for the 100 excitatory neurons in heterogeneous CA3 networks for E:I ratio = 100:10 (*A*), 100:20 (*B*), 100:50 (*C*). The columns represent the low, medium, and high trial-to-trial variability introduced by AGWN. *σ*_*noise*_ values for *Low*: 1 × 10^−6^ Hz^2^, *Medium*: 1 × 10^−2^ Hz^2^, *High*: 1 Hz^2^.

We then calculated the *SSI* metrics for all the noise levels and synaptic inhibition strengths and plotted the profiles of spatial information transfer for each level of E:I balance (Fig. 5). We observed heterogeneities in spatial information transfer profiles across different neurons in the heterogeneous network. Importantly, akin to the homogeneous networks, maximal information transfer occurred at the high-slope regions of the spatial tuning curves for low degrees of trial-to-trial variability, but transitioned to peak-firing location of the spatial tuning curves for high degrees of trial-to-trial variability (Fig. 5). Observations on heterogeneities and the consistent and significant reduction in both SSI metrics with increasing trial-to-trial variability were also noted from their quantifications (Fig. 6). Notably, similar to our observations with the homogeneous network (Fig. 3), there was no strong trend in *SSI peak* with increasing synaptic inhibition in heterogeneous networks as well (Fig. 6). *SSI FWHM* reduced significantly with increase in the strength of synaptic inhibition in both homogeneous (Fig. 3) and heterogeneous networks (Fig. 6).

**Figure 6:**
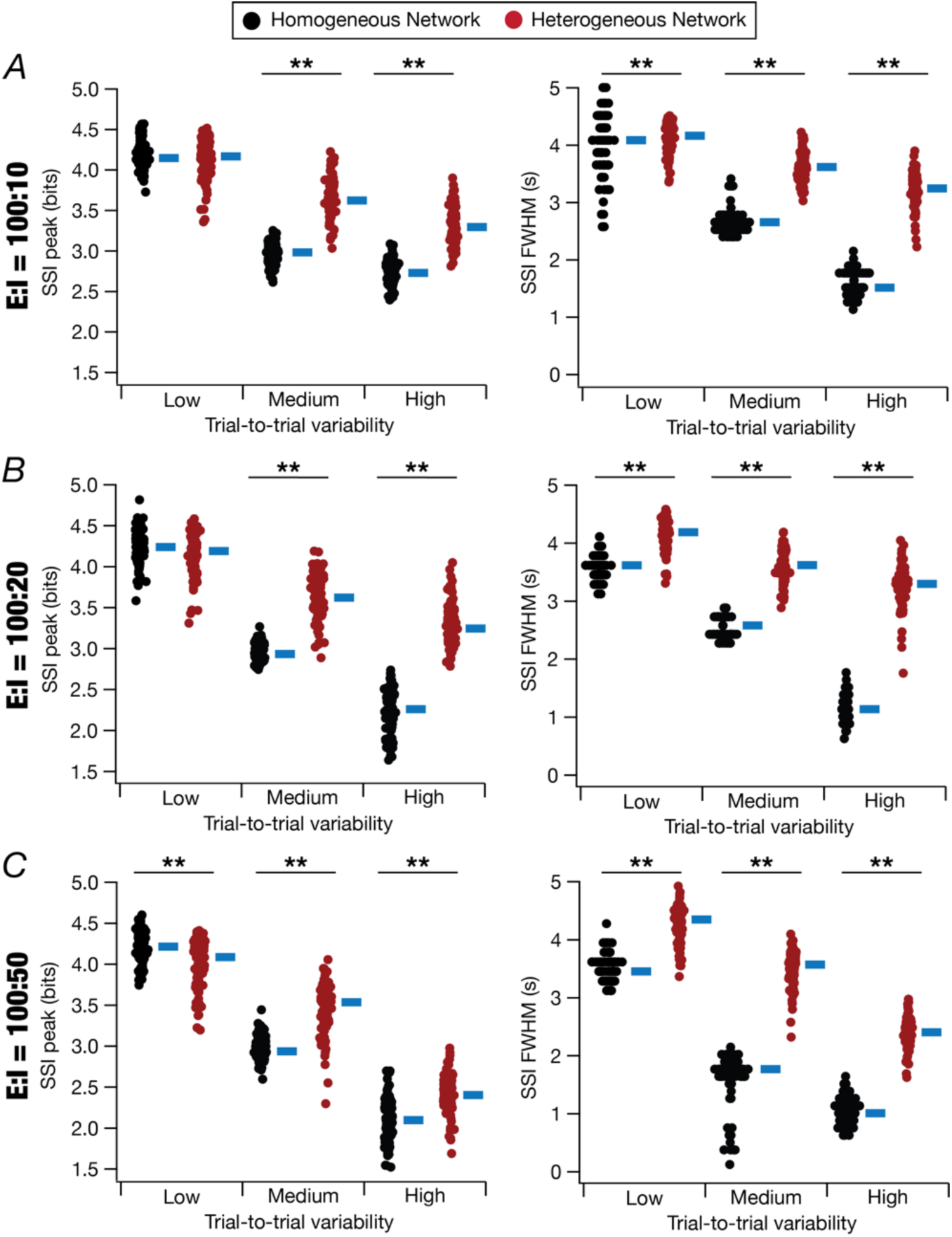
Spatial information transfer in heterogeneous networks manifested higher resilience to enhanced trial-to-trial variability compared to homogeneous networks. (*A*–*C*) Stimulus specific information (SSI) value at the center of the place field (*T*=5 s), *SSI peak*, and the full width at half-maxima, *SSI FWHM*, were computed to quantify the impact of trial-to-trial variability on the spatial information transfer across different synaptic E:I balances. Shown are these metrics for E:I balance at 100:10 (*A*), 100:20 (*B*), and 100: 50 (*C*). *σ*_*noise*_ values for *Low*: 1 × 10^−6^ Hz^2^, *Medium*: 1 × 10^−2^ Hz^2^, *High*: 1 Hz^2^. Outcomes of statistical tests: Panel *A,* For heterogeneous networks: *SSI peak*: Wilcoxon Rank Sum test, Low *vs*. Medium *p* = 2.2 × 10^−16^, Medium *vs*. High *p* = 2.2 × 10^−16^, Low *vs*. High *p* = 2.2 × 10^−16^, *SSI FWHM*: Wilcoxon Rank Sum test, Low *vs*. Medium *p* = 4.98 x 10^-14^, Medium *vs*. High *p* = 2.2 × 10^−16^, Low *vs*. High *p* = 2.2 × 10^−16^. Homogeneous *vs*. heterogeneous networks: *SSI peak*: Wilcoxon Rank Sum test, Low: *p* = 0.95, Medium: *p* = 2.2 × 10^−16^, High: *p* = 2.2 × 10^−16^. *SSI FWHM*: Wilcoxon Rank Sum test, Low: *p* = 2.2 × 10^−16^, Medium: *p* = 2.2 × 10^−16^, High: *p* = 2.2 × 10^−16^. Panel *B,* For heterogeneous networks, *SSI peak*: Wilcoxon Rank Sum test, Low *vs*. Medium *p* = 2.2 × 10^−16^, Medium *vs*. High *p* = 2.2 × 10^−16^, Low *vs*. High *p* = 2.2 × 10^−16^, *SSI FWHM*: Wilcoxon Rank Sum test, Low *vs*. Medium *p* = 2.2 × 10^−16^, Medium *vs*. High *p* = 2.2 × 10^−16^, Low *vs*. High *p* = 2.2 × 10^−16^. Homogeneous *vs*. heterogeneous networks: *SSI peak*: Wilcoxon Rank Sum test, Low: *p* = 0.12, Medium: *p* = 2.2 × 10^−16^, High: *p* = 2.2 × 10^−16^, *SSI FWHM*: Wilcoxon Rank Sum test, Low: *p* = 2.2 × 10^−16^, Medium: *p* = 2.2 × 10^−16^, High: *p* = 2.2 × 10^−16^. Panel *C,* For heterogeneous networks, *SSI peak*: Wilcoxon Rank Sum test, Low *vs*. Medium *p* = 2.2 × 10^−16^, Medium *vs*. High *p* = 2.4 × 10^−16^, Low *vs*. High *p* = 2.2 × 10^−16^, *SSI FWHM*: Wilcoxon Rank Sum test, Low *vs*. Medium *p* = 2.2 × 10^−16^, Medium *vs*. High *p* = 1.3 × 10^−11^, Low *vs*. High *p* = 2.2 × 10^−16^. Homogeneous *vs*. heterogeneous networks: *SSI peak*: Wilcoxon Rank Sum test, Low: *p* = 0.005, Medium: *p* = 2.2 × 10^−16^, High: *p* = 2.2 × 10^−16^, *SSI FWHM*: Wilcoxon Rank Sum test, Low: *p* = 2.2 × 10^−16^, Medium: *p* = 2.2 × 10^−16^, High: *p* = 2.2 × 10^−16^. The blue bars represent the respective median values for each of the measures.

Strikingly, we found significant increases in spatial information transfer in heterogeneous networks compared to their homogeneous counterparts, especially with higher degrees of trial-to-trial variability (Fig. 6). Specifically, while the *SSI peak* values were mostly comparable between homogeneous and heterogeneous networks for low degree of trial-to-trial variability, spatial information transfer was significantly more robust to increases in trial-to-trial variability in heterogeneous networks. Robustness of spatial information transfer to high trial-to-trial variability in heterogeneous networks was consistent across all levels of synaptic inhibition (Fig. 6). Quantitatively, the quantum of enhancement and the ranges of the *SSI peak* values were strongly dependent on the level of synaptic inhibition and the degree of trial-to-trial variability. In addition to the relative enhancement in the *SSI peak* values in heterogeneous networks compared to their homogeneous counterparts, we also found an increase in the *SSI FWHM*, especially in networks with high trial-to-trial variability and high synaptic inhibition (Fig. 6).

Together, spatial information transfer in heterogeneous networks was strikingly robust to high trial-to-trial variability and was strongly regulated by the level of synaptic inhibition and the degree of trial-to-trial variability.

### Spatial information transfer in the heterogeneous recurrent network for distinct spatially tuned inputs

Thus far, we analyzed the impact of synaptic E:I balance and trial-to-trial variability on spatial information transfer in intrinsically homogeneous (Figs. 1–3) and intrinsically heterogeneous (Figs. 4–6) networks, all receiving identically tuned afferent inputs. Despite the identical nature of spatial tuning of all afferent inputs impinging on the network, we found pronounced heterogeneities in firing rate and spatial information profiles in both homogeneous and heterogeneous networks. However, this single presynaptic place-field spike train does not account for heterogeneities associated with the tuning curves of the presynaptic neurons and differential synaptic weights associated with these presynaptic neurons. To account for this physiologically relevant scenario, we designed neuronal input structure such that they received stochastic inputs from all spatial locations, but acted as place cells for one location because of the differential synaptic weights (Bittner, et al., 2017; Dong, et al., 2021; D. Lee, Lin, & Lee, 2012; A. Roy & Narayanan, 2021; Seenivasan & Narayanan, 2020). We set the synaptic weight distribution such that spatially tuned responses of individual neurons occurred at distinct locations in the environment.

We simulated heterogeneous networks receiving such distinct place field inputs to elicit spatially tuned responses in each of the 100 excitatory neurons (Fig. 7). Neurons in these heterogeneous networks (endowed with afferent heterogeneities as well) showed place-field firing at distinct locations (Fig. 7*A*) with pronounced neuron-to-neuron variability in peak firing rate (Fig. 7*B*) and width (Fig. 7*C*) of the place-field firing profiles. While the peak firing rates and *FWHM* of networks with identical *vs*. distinct place-field centers broadly spanned similar ranges (Fig. 7*B–C*), there were significant increases in the peak firing frequencies of networks comprising neurons with distinct place-field centers across all E:I strengths (Fig. 7*B*). In addition, the median width of the place-field firing profiles as well as the neuron-to-neuron variability were higher in networks built of neurons with distinct place-field centers across all E:I strengths (Fig. 7*C*).

**Figure 7:**
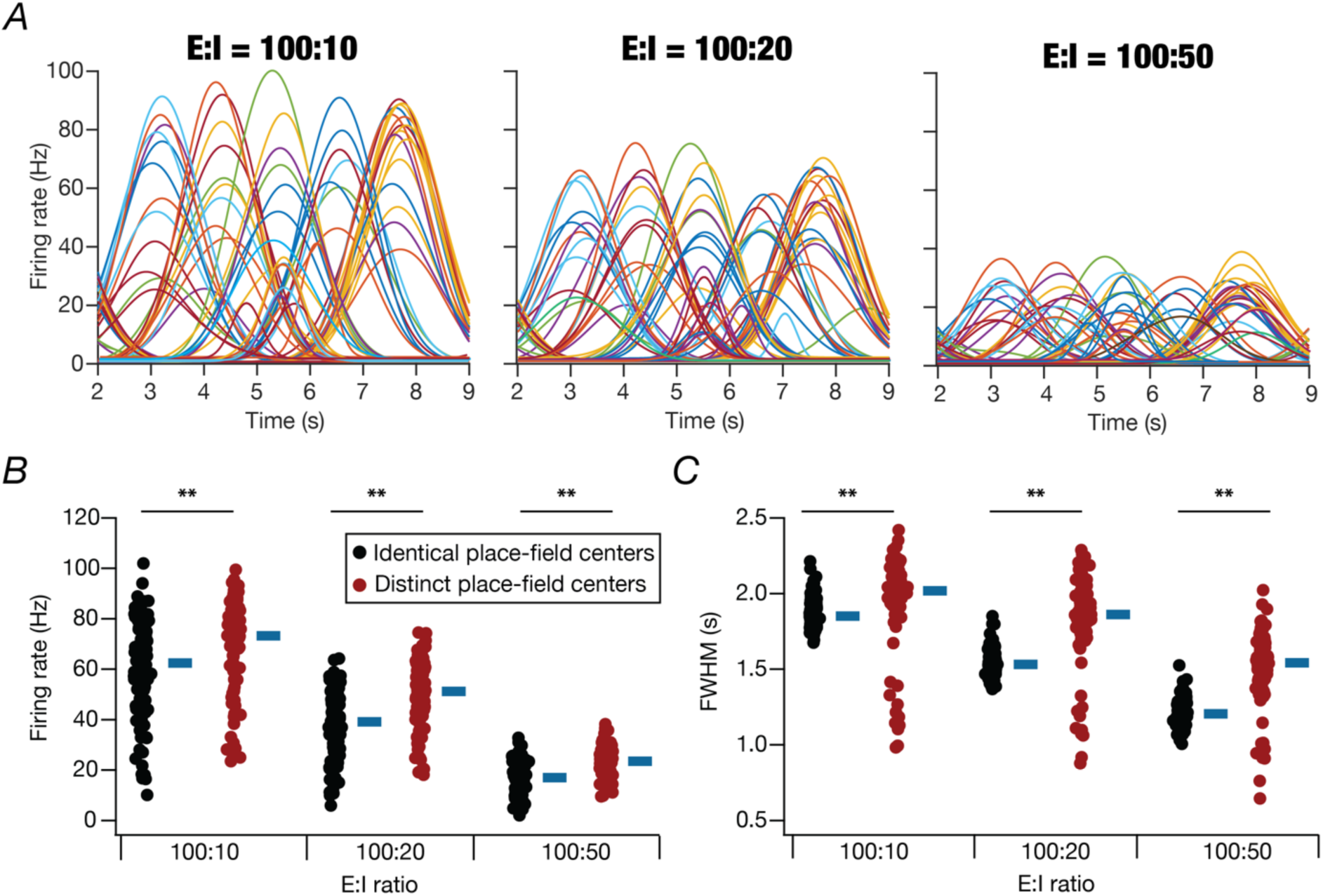
Place-field firing characteristics in heterogeneous networks with the distinct place-field centers. (*A*) Spatially tuned response of each excitatory neuron in the heterogeneous network with distinct place-field rcenters, plotted for all three cases of the E:I ratios. (*B*–*C*) Impact of different synaptic E:I strengths on the peak firing rate frequency, *F_max_* (*B*), and full-width at half maximum, *FWHM* of the firing rate profile (*C*) for each of the 100 excitatory neurons in the heterogeneous network. Shown are the comparisons with the firing rate and *FWHM* for neurons in networks with identical *vs.* distinct place-field centers. Statistical comparisons for *F_max_* in networks with identical *vs.* distinct place-field centers: Wilcoxon Rank Sum test, 100:10, *p* = 1.3 × 10^−3^, 100:20, *p* = 6.6 × 10^−9^, 100:50, *p* = 4.3 × 10^−10^. Statistical comparisons for *FWHM* in networks with identical *vs.* distinct place-field centers: Wilcoxon Rank Sum test, 100:10, *p* = 6.2 × 10^−10^, 100:20, *p* = 2.2 × 10^−16^, 100:50, p = 2.2 × 10^−16^.

We then performed 25 independent trials for 3 different levels of noise as well as 3 synaptic E:I strengths to compute the SSI tuning curve for each of the 100 spatially tuned neurons in this network. We quantified the spatial information transfer in each of the neurons in the heterogeneous network by computing the *SSI peak* at distinct spatial locations and the *SSI FWHM* of the corresponding spatial tuning curve (Fig. 8). We observed pronounced heterogeneities in the SSI measures across the network, with *SSI peak* reducing significantly with enhanced trial-to-trial variability and with increasing inhibitory strengths (Fig. 8). Importantly, we found a significant reduction in the *SSI peak* values for the network with distinct place-field inputs compared to those with identical place-field inputs. This was consistent across all degrees of trial-to-trial variability and all levels of synaptic inhibition. The dependence of *SSI FWHM* on identical *vs*. distinct place-field inputs was variable and was dependent on trial-to-trial variability. Whereas *SSI FWHM* was higher for the identical place-field inputs when trial-to-trial variability was low, the *SSI FWHM* for distinct place-field inputs was higher with increase in trial-to-trial variability. In most scenarios, the heterogeneity in both *SSI peak* and *SSI FWHM* values was higher in the network with distinct place-field inputs compared to those with identical place-field inputs (Fig. 8).

**Figure 8:**
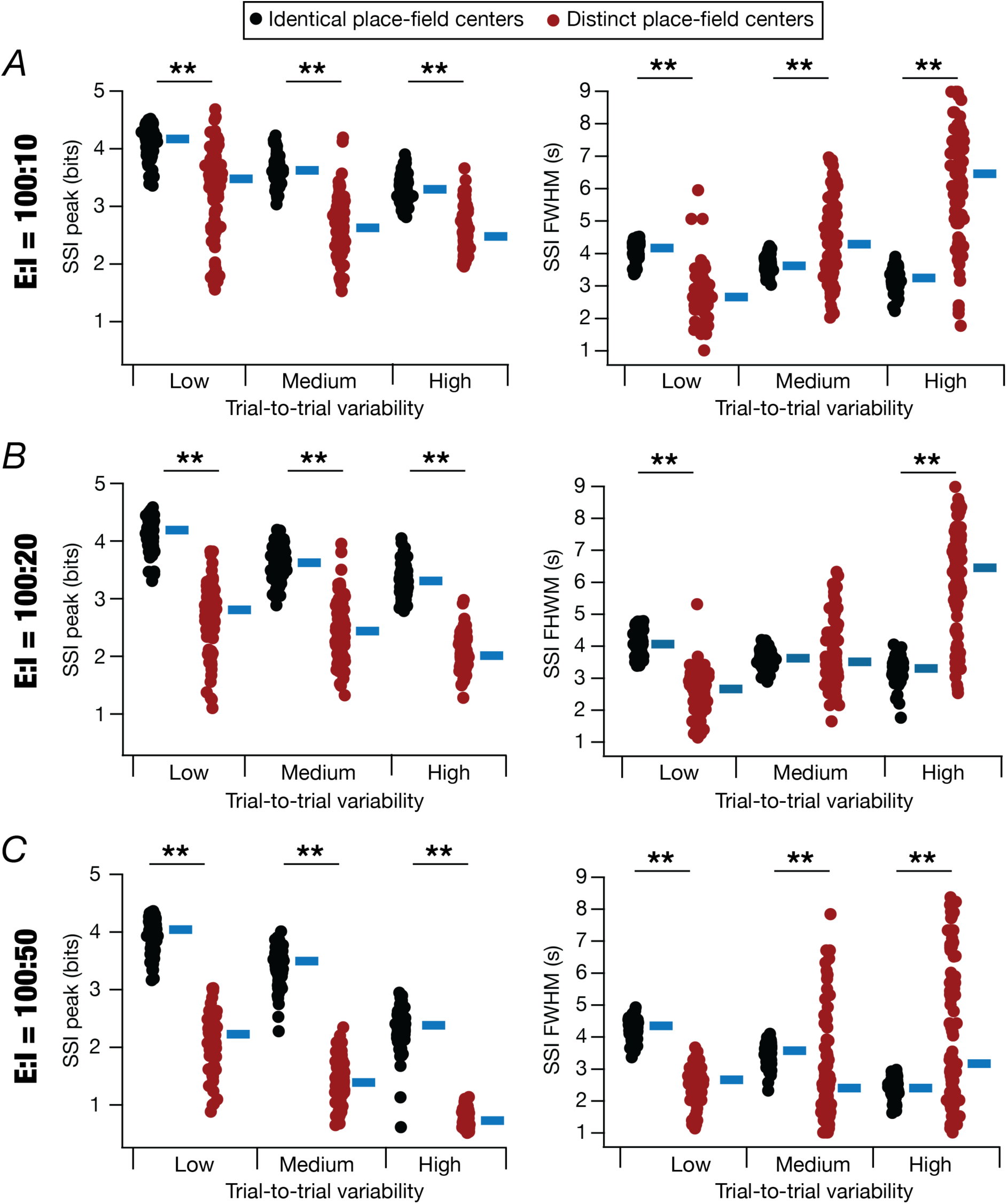
Spatial information transfer in heterogeneous networks was dependent on heterogeneities in place-field centers, synaptic inhibition, and trial-to-trial variability. (*A*–*C*) stimulus specific information (SSI) value at the distinct peak firing locations for each neuron in the heterogeneous network, *SSI peak*, and the full width at half-maxima, *SSI FWHM*, were computed to quantify the impact of trial-to-trial variability on the spatial information transfer across different values of E:I balance. Shown are metrics for heterogeneous networks with identical (Fig. 5) or distinct (Fig. 7) place-field centers. (*A*) *SSI peak* and *SSI FWHM* computed for E:I balance at 100:10, *SSI peak*: Kruskal Wallis test results for networks with distinct place-field centers, *p* = 0.481, Wilcoxon Rank Sum test results for networks with distinct place-field centers, Low *vs*. Medium *p* = 2 × 10^−13^, Medium *vs*. High *p* = 0.03, Low *vs*. High *p* = 5.7 × 10^−16^, *SSI FWHM*: Kruskal Wallis test results for networks with distinct place-field centers, *p* = 0.627, Wilcoxon Rank Sum test results for networks with distinct place-field centers, Low *vs*. Medium *p* = 4.98 × 10^−14^, Medium *vs*. High *p* = 3.5 × 10^−13^, Low *vs*. High *p* = 2.2 × 10^−16^. Statistical tests for networks with identical *vs*. distinct place-field centers: *SSI peak*: Wilcox Rank Sum test, Low: *p =* 2.2 × 10^−16^, Medium: *p* = 2.2 × 10^−16^, High: *p* = 2.2 × 10^−16^, *SSI FWHM* Low: *p =* 2.2 × 10^−16^, Medium: *p* = 1.13 × 10^−4^, High: *p* = 2.2 × 10^−16^. (*B*) *SSI peak* and *SSI FWHM* computed for E:I balance at 100:20, *SSI peak*: Kruskal Wallis test results for networks with distinct place-field centers, *p* = 0.481, Wilcoxon Rank Sum test results for networks with distinct place-field centers, Low *vs*. Medium *p* = 2 × 10^−5^, Medium *vs*. High *p* = 1 × 10^−10^, Low *vs*. High *p* = 2.2 × 10^−16^, *SSI FWHM*: Kruskal Wallis test results for networks with distinct place-field centers, *p* = 0.57, Wilcoxon Rank Sum test results for networks with distinct place-field centers, Low *vs*. Medium *p* = 2.2 × 10^−16^, Medium *vs*. High *p* = 7.8 × 10^−14^, Low *vs*. High *p* = 2.2 × 10^−16^. Statistical tests for networks with identical *vs*. distinct place-field centers: *SSI peak*: Wilcox Rank Sum test, Low: *p =* 2.2 × 10^−16^, Medium: *p* = 2.2 × 10^−16^, High: *p* = 2.2 × 10^−16^, *SSI FWHM* Low: *p =* 2.2 × 10^−16^, Medium: *p* = 0.3601, High: *p* = 2.2 × 10^−16^. (*C*) *SSI peak* and *SSI FWHM* computed for E:I balance at 100:50, *SSI peak*: Kruskal Wallis test results for networks with distinct place-field centers, *p* = 0.481, Wilcoxon Rank Sum test results for networks with distinct place-field centers, Low *vs*. Medium *p* = 2.2 × 10^−16^, Medium *vs*. High *p* = 2.2 × 10^−16^, Low *vs*. High *p* = 2.2 × 10^−16^, *SSI FWHM*: Kruskal Wallis test results for networks with distinct place-field centers, *p* = 0.146, Wilcoxon Rank Sum test results for networks with distinct place-field centers, Low *vs*. Medium *p* =0.006, Medium *vs*. High *p* = 0.02, Low *vs*. High *p* = 0.885. Statistical tests for networks with identical *vs*. distinct place-field centers: *SSI peak*: Wilcox Rank Sum test, Low: *p =* 2.2 × 10^−16^, Medium: *p* = 2.2 × 10^−16^, High: *p* = 2.2 × 10^−16^, *SSI FWHM* Low: *p =* 2.2 × 10^−16^, Medium: *p* = 1.4 × 10^−10^, High: *p* = 1.3 × 10^−4^. *σ*_*noise*_ values for network with distinct place-field centers *Low*: 1 × 10^−8^ Hz^2^, *Medium*: 1 × 10^−6^ Hz^2^, *High*: 1 × 10^−5^ Hz^2^. The blue bars represent the respective median values for each of the measures.

Together, our analyses demonstrated that the efficacy of spatial information transfer across neurons in recurrent networks was regulated by several network components: excitation-inhibition balance, trial-to-trial variability, intrinsic neuronal heterogeneities, and afferent heterogeneities (Figs. 1–8). These observations also highlight the critical roles of synaptic wiring and neuronal intrinsic properties in the emergence of spatial tuning and in the effectiveness of spatial information transfer in excitatory neurons of recurrent spiking networks.

## DISCUSSION

We demonstrated pronounced heterogeneities in spatial tuning profiles and spatial information transfer across place cells within recurrent spiking networks. First, we unveiled pronounced heterogeneities in the peak firing rate, the width of place-field, subthreshold voltage ramp, and spatial information transfer in different neurons even within networks composed of homogeneous repeating units that received identically tuned spatial inputs. Importantly, we found a strong dependence of the spatial tuning profiles and spatial information transfer across neurons in these homogeneous networks on the degree of trial-to-trial variability. Specifically, consistent with earlier observations with morphologically realistic single-neuron models of CA1 place cells (A. Roy & Narayanan, 2021), we found that increasing the degree of trial-to-trial variability resulted in a reduction in spatial information transfer. In addition, with increasing trial-to-trial variability, the spatial location of peak information transfer transitioned from high slope regions to the peak region of the corresponding spatial tuning curve. Although enhanced inhibition expectedly resulted in reductions in firing rate, there was no major impact of inhibitory balance on the peak value of spatial information transferred. However, there were reductions in both the extent of place-field firing and the width of the spatial information transfer profile with increase in inhibition. These observations reveal heterogeneities in spatial information transfer even in a homogeneous network, with strong dependence of the information transfer on trial-to-trial variability.

We then introduced within-type intrinsic heterogeneities into both excitatory and inhibitory neurons of the recurrent network and presented identically tuned spatial inputs to these neurons. In these heterogeneous recurrent networks, the strength of synaptic inhibition played a critical role in regulating the extent of heterogeneities in spatial tuning profiles and highlighted the differences in heterogeneous networks from their homogeneous counterparts. Strikingly, spatial information transfer in heterogeneous networks was robust to high trial-to-trial variability when compared to their homogeneous counterparts. Spatial information transfer was also strongly regulated by the level of synaptic inhibition and the degree of trial-to-trial variability in these heterogeneous networks. Finally, we introduced afferent heterogeneities into these intrinsically heterogeneous networks by providing distinct synaptic inputs to individual neurons, which allowed them to be spatially selective to different locations. Afferent heterogeneities introduced a significant reduction in the peak information transfer values for neurons in the network with distinct place-field inputs compared to those with identical place-field inputs. In addition, afferent heterogeneities also enhanced the extent of heterogeneities in spatial information transfer profiles across neurons in the network.

Together, our analyses demonstrated that the efficacy of spatial information transfer across neurons in recurrent networks was regulated by several network components: excitation-inhibition balance, trial-to-trial variability, intrinsic neuronal heterogeneities, and afferent heterogeneities. These observations also highlight the critical roles of synaptic wiring and neuronal intrinsic properties in the emergence of spatial tuning and in the efficacy of spatial information transfer in excitatory neurons of recurrent spiking networks. Importantly, with several disparate routes (involving a diversity of neuronal and circuit components) to achieving similar spatial information transfer, our study highlights the manifestation of degeneracy in the emergence of strong spatial tuning and efficacious spatial information transfer in such networks. Finally, our results also underscore the critical role of within-type intrinsic neuronal heterogeneities in enhancing the functional robustness of spatial information transfer in the face of perturbations.

### Degeneracy in and robustness of spatial information transfer in homogeneous and heterogeneous networks

Spatial tuning and spatial information transfer across individual hippocampal pyramidal neurons have been reported to manifest cellular-scale degeneracy (Basak & Narayanan, 2018, 2020; A. Roy & Narayanan, 2021; Seenivasan & Narayanan, 2020), as have other intrinsic properties in neurons of the hippocampal formation (Kumari & Narayanan, 2024; Mishra & Narayanan, 2019, 2021a; Mittal & Narayanan, 2018, 2022; Rathour, Malik, & Narayanan, 2016; Rathour & Narayanan, 2014; R. Roy & Narayanan, 2022). Specifically, these analyses showed that disparate morphological and molecular (*e.g.*, ion channels, buffers) components could collectively yield similar spatial tuning curves (Basak & Narayanan, 2018, 2020; A. Roy & Narayanan, 2021; Seenivasan & Narayanan, 2020) and similar spatial information transfer through rate (A. Roy & Narayanan, 2021) or phase (Seenivasan & Narayanan, 2020) codes of space.

Through systematic evaluation of the impact of different neuronal and circuit components, we demonstrated that similar tuning curves and spatial information transfer profiles of neurons in a recurrent network could be achieved through disparate combinations of different components. We observed variable dependencies of spatial tuning profiles and the associated spatial information transfer profiles to specific circuit components. For instance, in homogeneous networks, while inhibition had a strong role to play in regulating peak firing rate (Fig. 1*C*), *SSI peak* was largely unaffected (Fig. 3) by changes in the synaptic E:I balance. Thus, although different measurements were dependent on disparate set of parametric combinations, there were several components that contributed to each measurement. For instance, peak spatial information transfer was critically dependent on intrinsic heterogeneities, trial-to-trial variability, and afferent heterogeneities. In addition, any given component affected several functional measurements. As an example, trial-to-trial variability affected peak firing rates, firing rate profiles, peak SSI, and the shape of spatial information transfer. Together, these observations point to a many-to-many mapping between specific attributes and functional measurements, with strong interdependencies across each other. Such many-to-many mappings are hallmarks of systems manifesting degeneracy (the ability of disparate components to yield similar functional outcomes) and pleiotropy (the ability of the same components to contribute to different functional measurements in a context dependent manner) (Albantakis, Bernard, Brenner, Marder, & Narayanan, 2024; Calabrese & Marder, 2025; Edelman & Gally, 2001; Mishra & Narayanan, 2019, 2021b; Rathour & Narayanan, 2019; J. Yang & Prescott, 2023).

Apart from demonstrating the manifestation of degeneracy, our analyses also emphasize the role of within-type heterogeneities in neural circuit function, specifically with reference to spatial information transfer. Specifically, our analyses show that intrinsically heterogeneous networks were more robust to perturbations compared to their homogeneous counterparts (Fig. 6). Within-type heterogeneities are ubiquitous (Cembrowski & Spruston, 2019; Mishra & Narayanan, 2020; Planert, et al., 2025; R. Roy & Narayanan, 2023; Sun, Jiang, & Lu, 2020; Sun, et al., 2017). However, the impact of within-type heterogeneities on neural-circuit function has been shown to be diverse, depending on several factors including the form of heterogeneities, degree of heterogeneities, the task under consideration, the performance metrics used, the specific models employed, the network architecture, and interactions among different forms of heterogeneities (Dahmen, et al., 2025; Gast, et al., 2024; Mishra & Narayanan, 2019, 2021b; Mittal & Narayanan, 2021; Rich, Moradi Chameh, Lefebvre, & Valiante, 2022; Saini & Narayanan, 2025; Santhosh & Narayanan, 2025). In our case, we studied the impact of synaptic, intrinsic, and afferent heterogeneities on spatial information transfer in recurrent networks to demonstrate the dependence on the specific form of heterogeneity. Whereas afferent synaptic heterogeneities translated to heterogeneities in spatial information transfer, changes to local-synaptic E:I balance altered information transfer profiles. Intrinsic and afferent heterogeneities also interacted with synaptic heterogeneities and trial-to-trial variability in complex ways towards modulating spatial information transfer. Together, these observations further emphasize the need to explicitly account for different forms of heterogeneities and their interactions with each other in studying the impact of heterogeneities on neural-circuit function.

### Impact of trial-to-trial variability on spatial information transfer in homogeneous and heterogeneous networks

Our results discern the debilitating impact of trial-to-trial variability on spatial information transfer across neurons in a recurrent CA3 network model. With increase in trial-to-trial variability, we found a significant reduction in spatial information transfer across neurons in both the homogeneous and heterogeneous networks, in a manner that was consistent with prior observations from a heterogeneous population of individual CA1 pyramidal neuron models (A. Roy & Narayanan, 2021). These observations imply that physiological or pathological conditions that enhance trial-to-trial variability would be detrimental to information transfer in recurrent networks.

Our study also extends earlier findings about the relationship between tuning curves and stimulus-specific spatial information to a network context. Specifically, earlier results spanning different tuning curves have shown that SSI profiles transitioned from bimodal to unimodal distributions with increase in trial-to-trial variability (Butts & Goldman, 2006; Montgomery & Wehr, 2010; A. Roy & Narayanan, 2021). This transition was consequent to a shift in the peak of the SSI profile from high slope regions of the tuning curve during low trial-to-trial variability to high-response regions of the tuning curve with high trial-to-trial variability. Our findings with spatial information transfer are consistent with these observations, across both homogeneous and heterogeneous networks. Together, our observations highlight the critical need to account for neural-circuit heterogeneities, trial-to-trial variability, and the intricate interactions among them in assessing neural tuning curves as well as information transfer.

### Inhibitory control of spatial information transfer in the homogeneous and heterogeneous networks

A prominent source of heterogeneities in the network physiology emerges from differences in synaptic strength values, not just from afferent synaptic connectivity but also local connectivity. In a recurrent network, where there is feedback excitation, it is essential to have sources of inhibitory inputs stabilize the network and ensure that there is no runaway excitation (Brunel, 2000; Sprekeler, 2017; Van Vreeswijk & Sompolinsky, 1996). As afferent and recurrent feedback synaptic strengths could be heterogeneous across different neurons, granular forms of excitation-inhibition balance also manifest as heterogeneities in inhibitory synaptic weights. Thus heterogeneities in excitatory and inhibitory inputs and balance between them form a crucial component for the maintenance of stability and homeostasis in neuronal circuits (Bhatia, et al., 2019; Mishra & Narayanan, 2021c; Rubin, Abbott, & Sompolinsky, 2017; Sukenik, et al., 2021; Taub, Katz, & Lampl, 2013; W. Yang & Sun, 2018; Yu, et al., 2018; S. Zhou & Y. Yu, 2018).

Inhibition has also been strongly implicated in shaping place field responses in hippocampal place cells (Grienberger, et al., 2017). One of the mechanisms by which synaptic inhibition could possibly shape the place cell responses is by suppressing the off-target and out-of-field responses. We observed that increasing synaptic inhibition resulted in reduction of place field firing, width of the spatial tuning profile, and the characteristic ramp voltage. However, the impact of inhibition on peak information transfer was minimal if the afferent inputs were homogeneous, and irrespective of whether the intrinsic properties of the network were homogeneous (Fig. 3) or heterogeneous (Fig. 6). However, with the introduction of afferent heterogeneities, increasing inhibition reduced spatial information transfer as well (Fig. 8). This is to be expected because inhibitory neurons will be active across several locations in networks receiving heterogeneous afferent inputs. Such spatially widespread activation of inhibition would therefore suppress activity of several cells, together contributing to reduced spatial information transfer. These observations emphasize the strong interactions between local and afferent synaptic heterogeneities, showing the dependence of measured function on cross-interactions between both forms of heterogeneities. Together, our results highlight the critical role of E:I balance in neural circuits, also underscoring the need to assess the impact of all forms of heterogeneities together, rather than studying heterogeneities in individual components in a piecemeal fashion.

### Limitations and future directions

We introduced intrinsic heterogeneities by assigning different values to each parameter from a random uniform distribution such that the excitatory and the inhibitory neurons exhibited tonic-bursting profiles and fast-spiking profiles respectively. The emergence of sharply-tuned responses in place cells has been attributed to a plethora of mechanisms such as disparate non-random combinations of molecular components (Basak & Narayanan, 2018, 2020; A. Roy & Narayanan, 2021; Seenivasan & Narayanan, 2020), spatial dispersion of the excitatory synapses along the somato-dendritic axis (Basak & Narayanan, 2018, 2020; A. Roy & Narayanan, 2021), morphological attributes (Basak & Narayanan, 2020), dendritic spikes and plateau potentials (Basak & Narayanan, 2018; Bittner, et al., 2015; Gasparini, Migliore, & Magee, 2004; Golding, Jung, Mickus, & Spruston, 1999; Golding & Spruston, 1998; Losonczy & Magee, 2006; Royer, et al., 2012; M. E. Sheffield & Dombeck, 2015; M. E. J. Sheffield, Adoff, & Dombeck, 2017), behavioral relevance of spatial locations (Dupret, O’Neill, Pleydell-Bouverie, & Csicsvari, 2010; Gauthier & Tank, 2018; H. Lee, Ghim, Kim, Lee, & Jung, 2012; Michon, Krul, Sun, & Kloosterman, 2021; Robinson, et al., 2020; Sosa, Plitt, & Giocomo, 2025), preexisting network dynamics (McKenzie, et al., 2021; Zutshi, Valero, Fernandez-Ruiz, & Buzsaki, 2022), and different sources of inhibition (Campbell, Martin, Magee, & Grienberger, 2025; Fernandez-Arroyo, Jurado, & Lerma, 2025; Grienberger, et al., 2017; Royer, et al., 2012).

Since the parameters in the Izhikevich models only account for the spiking properties, the parametric space here does not account for detailed biophysics, morphological characteristics, dendritic physiology, or reward-modulation of biophysical mechanisms. Future studies should be directed towards exploring the spatial information transfer in a recurrent network comprised of morphologically and biophysically realistic CA3 neurons where biophysical mechanisms and reward modulation can be probed in detail. Spatio-temporal dynamics of the excitation-inhibition balance in individual neurons within the recurrent network can also be studied to understand how the precise excitation-inhibition balance (Bhatia, et al., 2019; Campbell, et al., 2025; Fernandez-Arroyo, et al., 2025; Grienberger, et al., 2017; Royer, et al., 2012; Sprekeler, 2017; Yu, et al., 2018) can regulate the spatial information transfer in heterogeneous CA3 recurrent networks. Such networks built of neurons that account for morphology could also account for strata-specific inputs onto CA3 pyramidal neurons through the mossy fibers from the dentate gyrus, the commissural/associational fibers from CA3, and the perforant pathway from the entorhinal cortex (P. Andersen, et al., 2006). Finally, in assessing trial-to-trial variability, different colors of channel noise *vs*. synaptic noise (Faisal, Selen, & Wolpert, 2008) could be studied, instead of our limited use of additive Gaussian white noise in our analysis.

## Author Contributions

R. R. and R. N. designed experiments; R. R. performed experiments; R. R. analyzed data; R. R. and R. N. wrote the paper.

## Competing Interest Statement

The authors declare that they have no competing interests.

## Acknowledgments

The authors thank members of the cellular neurophysiology laboratory for helpful discussions and for comments on a draft of this manuscript. This work was supported by the Ministry of education (R. R. and R. N.).

## Data Availability Statement

All data required to evaluate this manuscript is available as part of the manuscript.

## Notes

### Competing Interest Statement

The authors have declared no competing interest.

## REFERENCES

Albantakis, L., Bernard, C., Brenner, N., Marder, E., & Narayanan, R. (2024). The brain’s best kept secret is its degenerate structure. J Neurosci, 44.

Andersen, P., Morris, R., Amaral, D., Bliss, T., & O’Keefe, J. (2006). The hippocampus book. New York, USA: Oxford University Press.

Andersen, P., Morris, R., Amaral, D., Bliss, T., & O’Keefe, J. (2007). *The hippocampus book*: Oxford university press.

Andrews, B. W., & Iglesias, P. A. (2007). An information-theoretic characterization of the optimal gradient sensing response of cells. PLoS computational biology, 3, e153.

Attneave, F. (1954). Some informational aspects of visual perception. Psychol Rev, 61, 183–193.

Barlow, H. (1961). Possible principles underlying the transformation of sensory messages. In. WA Rosenblith (Ed.), Sensory communications. In: Cambridge, MA: MIT Press.

Basak, R., & Narayanan, R. (2018). Spatially dispersed synapses yield sharply-tuned place cell responses through dendritic spike initiation. Journal of Physiology, 596, 4173–4205.

Basak, R., & Narayanan, R. (2020). Robust emergence of sharply tuned place-cell responses in hippocampal neurons with structural and biophysical heterogeneities. Brain Struct Funct, 225, 567–590.

Bell, A. J., & Sejnowski, T. J. (1997). The “independent components” of natural scenes are edge filters. Vision research, 37, 3327–3338.

Bennett, M. R., Gibson, W. G., & Robinson, J. (1994). Dynamics of the CA3 pyramidal neuron autoassociative memory network in the hippocampus. Philos Trans R Soc Lond B Biol Sci, 343, 167–187.

Bhatia, A., Moza, S., & Bhalla, U. S. (2019). Precise excitation-inhibition balance controls gain and timing in the hippocampus. Elife, 8.

Bittner, K. C., Grienberger, C., Vaidya, S. P., Milstein, A. D., Macklin, J. J., Suh, J., Tonegawa, S., & Magee, J. C. (2015). Conjunctive input processing drives feature selectivity in hippocampal CA1 neurons. Nat Neurosci, 18, 1133–1142.

Bittner, K. C., Milstein, A. D., Grienberger, C., Romani, S., & Magee, J. C. (2017). Behavioral time scale synaptic plasticity underlies CA1 place fields. Science, 357, 1033–1036.

Brenner, N., Bialek, W., & Van Steveninck, R. d. R. (2000). Adaptive rescaling maximizes information transmission. Neuron, 26, 695–702.

Brunel, N. (2000). Dynamics of sparsely connected networks of excitatory and inhibitory spiking neurons. Journal of computational neuroscience, 8, 183–208.

Butts, D. A. (2003). How much information is associated with a particular stimulus? Network, 14, 177–187.

Butts, D. A., & Goldman, M. S. (2006). Tuning curves, neuronal variability, and sensory coding. PLoS Biol, 4, e92.

Buzsaki, G., & Moser, E. I. (2013). Memory, navigation and theta rhythm in the hippocampal-entorhinal system. Nat Neurosci, 16, 130–138.

Calabrese, R. L., & Marder, E. (2025). Degenerate neuronal and circuit mechanisms important for generating rhythmic motor patterns. Physiol Rev, 105, 95–135.

Campbell, E. P., Martin, L., Magee, J. C., & Grienberger, C. (2025). Dendrite-targeting OLM interneurons regulate the formation of learning-related CA1 place cell representations. bioRxiv.

Cembrowski, M. S., & Spruston, N. (2019). Heterogeneity within classical cell types is the rule: lessons from hippocampal pyramidal neurons. Nat Rev Neurosci, 20, 193–204.

Dahmen, D., Hutt, A., Indiveri, G., Kennedy, A., Lefebvre, J., Mazzucato, L., Motter, A. E., Narayanan, R., Payvand, M., Planert, H., & Gast, R. (2025). How Heterogeneity Shapes Dynamics and Computation in the Brain. Neuron, *In press*, DOI: 10.1016/j.neuron.2025.1011.1023.

DeWeese, M. R., & Meister, M. (1999). How to measure the information gained from one symbol. Network, 10, 325–340.

Dombeck, D. A., Harvey, C. D., Tian, L., Looger, L. L., & Tank, D. W. (2010). Functional imaging of hippocampal place cells at cellular resolution during virtual navigation. Nat Neurosci, 13, 1433–1440.

Dong, C., Madar, A. D., & Sheffield, M. E. J. (2021). Distinct place cell dynamics in CA1 and CA3 encode experience in new environments. Nat Commun, 12, 2977.

Dupret, D., O’Neill, J., Pleydell-Bouverie, B., & Csicsvari, J. (2010). The reorganization and reactivation of hippocampal maps predict spatial memory performance. Nat Neurosci, 13, 995–1002.

Edelman, G. M., & Gally, J. A. (2001). Degeneracy and complexity in biological systems. Proc Natl Acad Sci U S A, 98, 13763–13768.

Fairhall, A. L., Lewen, G. D., Bialek, W., & de Ruyter van Steveninck, R. R. (2001). Efficiency and ambiguity in an adaptive neural code. Nature, 412, 787–792.

Faisal, A. A., Selen, L. P., & Wolpert, D. M. (2008). Noise in the nervous system. Nat Rev Neurosci, 9, 292–303.

Fernandez-Arroyo, B., Jurado, S., & Lerma, J. (2025). Understanding OLM interneurons: Characterization, circuitry, and significance in memory and navigation. Neuroscience, 578, 69–80.

Gasparini, S., Migliore, M., & Magee, J. C. (2004). On the initiation and propagation of dendritic spikes in CA1 pyramidal neurons. J Neurosci, 24, 11046–11056.

Gast, R., Solla, S. A., & Kennedy, A. (2024). Neural heterogeneity controls computations in spiking neural networks. Proc Natl Acad Sci U S A, 121, e2311885121.

Gauthier, J. L., & Tank, D. W. (2018). A Dedicated Population for Reward Coding in the Hippocampus. Neuron, 99, 179–193 e177.

Geisler, C., Diba, K., Pastalkova, E., Mizuseki, K., Royer, S., & Buzsaki, G. (2010). Temporal delays among place cells determine the frequency of population theta oscillations in the hippocampus. Proc Natl Acad Sci U S A, 107, 7957–7962.

Gloveli, T., Dugladze, T., Saha, S., Monyer, H., Heinemann, U., Traub, R. D., Whittington, M. A., & Buhl, E. H. (2005). Differential involvement of oriens/pyramidale interneurones in hippocampal network oscillations in vitro. Journal of Physiology, 562, 131–147.

Golding, N. L., Jung, H. Y., Mickus, T., & Spruston, N. (1999). Dendritic calcium spike initiation and repolarization are controlled by distinct potassium channel subtypes in CA1 pyramidal neurons. J Neurosci, 19, 8789–8798.

Golding, N. L., & Spruston, N. (1998). Dendritic sodium spikes are variable triggers of axonal action potentials in hippocampal CA1 pyramidal neurons. Neuron, 21, 1189–1200.

Golomb, D., Donner, K., Shacham, L., Shlosberg, D., Amitai, Y., & Hansel, D. (2007). Mechanisms of firing patterns in fast-spiking cortical interneurons. PLoS Comput Biol, 3, e156.

Grienberger, C., Milstein, A. D., Bittner, K. C., Romani, S., & Magee, J. C. (2017). Inhibitory suppression of heterogeneously tuned excitation enhances spatial coding in CA1 place cells. Nat Neurosci, 20, 417–426.

Harvey, C. D., Collman, F., Dombeck, D. A., & Tank, D. W. (2009). Intracellular dynamics of hippocampal place cells during virtual navigation. Nature, 461, 941–946.

Hemond, P., Epstein, D., Boley, A., Migliore, M., Ascoli, G. A., & Jaffe, D. B. (2008). Distinct classes of pyramidal cells exhibit mutually exclusive firing patterns in hippocampal area CA3b. Hippocampus, 18, 411–424.

Hubel, D. H., & Wiesel, T. N. (1962). Receptive fields, binocular interaction and functional architecture in the cat’s visual cortex. J Physiol, 160, 106–154.

Izhikevich, E. M. (2003). Simple model of spiking neurons. IEEE Trans Neural Netw, 14, 1569–1572.

Káli, S., & Dayan, P. (2000). The Involvement of Recurrent Connections in Area CA3 in Establishing the Properties of Place Fields: a Model. The Journal of Neuroscience, 20, 7463–7477.

Kawaguchi, Y. (2001). Distinct firing patterns of neuronal subtypes in cortical synchronized activities. J Neurosci, 21, 7261–7272.

Kumari, S., & Narayanan, R. (2024). Ion-channel degeneracy and heterogeneities in the emergence of signature physiological characteristics of dentate gyrus granule cells. J Neurophysiol, 132, 991–1013.

Laughlin, S. (1981). A simple coding procedure enhances a neuron’s information capacity. Zeitschrift für Naturforschung c, 36, 910–912.

Le Duigou, C., Simonnet, J., Telenczuk, M. T., Fricker, D., & Miles, R. (2014). Recurrent synapses and circuits in the CA3 region of the hippocampus: an associative network. Front Cell Neurosci, 7, 262.

Lee, D., Lin, B. J., & Lee, A. K. (2012). Hippocampal place fields emerge upon single-cell manipulation of excitability during behavior. Science, 337, 849–853.

Lee, H., Ghim, J. W., Kim, H., Lee, D., & Jung, M. (2012). Hippocampal neural correlates for values of experienced events. J Neurosci, 32, 15053–15065.

Li, Y., Briguglio, J. J., Romani, S., & Magee, J. C. (2024). Mechanisms of memory-supporting neuronal dynamics in hippocampal area CA3. Cell, 187, 6804–6819 e6821.

Losonczy, A., & Magee, J. C. (2006). Integrative properties of radial oblique dendrites in hippocampal CA1 pyramidal neurons. Neuron, 50, 291–307.

Lundstrom, B. N., Higgs, M. H., Spain, W. J., & Fairhall, A. L. (2008). Fractional differentiation by neocortical pyramidal neurons. Nature neuroscience, 11, 1335–1342.

Mackwood, O., Naumann, L. B., & Sprekeler, H. (2021). Learning excitatory-inhibitory neuronal assemblies in recurrent networks. Elife, 10.

Marder, E., & Taylor, A. L. (2011). Multiple models to capture the variability in biological neurons and networks. Nat Neurosci, 14, 133–138.

Markram, H., Toledo-Rodriguez, M., Wang, Y., Gupta, A., Silberberg, G., & Wu, C. (2004). Interneurons of the neocortical inhibitory system. Nature Reviews Neuroscience, 5, 793–807.

McCormick, D. A., Connors, B. W., Lighthall, J. W., & Prince, D. A. (1985). Comparative electrophysiology of pyramidal and sparsely spiny stellate neurons of the neocortex. Journal of neurophysiology, 54, 782–806.

McKenzie, S., Huszár, R., English, D. F., Kim, K., Christensen, F., Yoon, E., & Buzsáki, G. (2021). Preexisting hippocampal network dynamics constrain optogenetically induced place fields. Neuron, 109, 1040–1054.e1047.

Michon, F., Krul, E., Sun, J. J., & Kloosterman, F. (2021). Single-trial dynamics of hippocampal spatial representations are modulated by reward value. Curr Biol, 31, 4423–4435 e4425.

Mishra, P., & Narayanan, R. (2019). Disparate forms of heterogeneities and interactions among them drive channel decorrelation in the dentate gyrus: Degeneracy and dominance. Hippocampus, 29, 378–403.

Mishra, P., & Narayanan, R. (2020). Heterogeneities in intrinsic excitability and frequency-dependent response properties of granule cells across the blades of the rat dentate gyrus. J Neurophysiol, 123, 755–772.

Mishra, P., & Narayanan, R. (2021a). Ion-channel degeneracy: Multiple ion channels heterogeneously regulate intrinsic physiology of rat hippocampal granule cells. Physiol Rep, 9, e14963.

Mishra, P., & Narayanan, R. (2021b). Ion-channel regulation of response decorrelation in a heterogeneous multi-scale model of the dentate gyrus. Curr Res Neurobiol, 2, 100007.

Mishra, P., & Narayanan, R. (2021c). Stable continual learning through structured multiscale plasticity manifolds. Curr Opin Neurobiol, 70, 51–63.

Mittal, D., & Narayanan, R. (2018). Degeneracy in the robust expression of spectral selectivity, subthreshold oscillations, and intrinsic excitability of entorhinal stellate cells. J Neurophysiol, 120, 576–600.

Mittal, D., & Narayanan, R. (2021). Resonating neurons stabilize heterogeneous grid-cell networks. Elife, 10, e66804.

Mittal, D., & Narayanan, R. (2022). Heterogeneous stochastic bifurcations explain intrinsic oscillatory patterns in entorhinal cortical stellate cells. Proc Natl Acad Sci U S A, 119, e2202962119.

Montgomery, N., & Wehr, M. (2010). Auditory cortical neurons convey maximal stimulus-specific information at their best frequency. J Neurosci, 30, 13362–13366.

Moser, E. I., Kropff, E., & Moser, M.-B. (2008). Place cells, grid cells, and the brain’s spatial representation system. Annu. Rev. Neurosci., 31, 69–89.

Moser, M. B., Rowland, D. C., & Moser, E. I. (2015). Place cells, grid cells, and memory. Cold Spring Harb Perspect Biol, 7, a021808.

Murphy, B. K., & Miller, K. D. (2009). Supplemental Materials for “Balanced amplification: a new mechanism of selective amplification of neural activity patterns”.

Narayanan, R., & Johnston, D. (2012). Functional maps within a single neuron. Journal of neurophysiology, 108, 2343–2351.

O’Keefe, J. (1976). Place units in the hippocampus of the freely moving rat. Experimental neurology, 51, 78–109.

O’Keefe, J., & Dostrovsky, J. (1971). The hippocampus as a spatial map. Preliminary evidence from unit activity in the freely-moving rat. Brain Res, 34, 171–175.

Panzeri, S., Senatore, R., Montemurro, M. A., & Petersen, R. S. (2007). Correcting for the sampling bias problem in spike train information measures. Journal of neurophysiology, 98, 1064–1072.

Planert, H., Mittermaier, F. X., Grosser, S., Fidzinski, P., Schneider, U. C., Radbruch, H., Onken, J., Holtkamp, M., Schmitz, D., Alle, H., Vida, I., Geiger, J. R. P., & Peng, Y. (2025). Electrophysiological classification of human layer 2-3 pyramidal neurons reveals subtype-specific synaptic interactions. Nat Neurosci.

Rathour, R. K., Malik, R., & Narayanan, R. (2016). Transient potassium channels augment degeneracy in hippocampal active dendritic spectral tuning. Sci Rep, 6, 24678.

Rathour, R. K., & Narayanan, R. (2014). Homeostasis of functional maps in active dendrites emerges in the absence of individual channelostasis. Proc Natl Acad Sci U S A, 111, E1787–1796.

Rathour, R. K., & Narayanan, R. (2019). Degeneracy in hippocampal physiology and plasticity. Hippocampus, 29, 980–1022.

Rich, S., Moradi Chameh, H., Lefebvre, J., & Valiante, T. A. (2022). Loss of neuronal heterogeneity in epileptogenic human tissue impairs network resilience to sudden changes in synchrony. Cell Rep, 39, 110863.

Robinson, N. T. M., Descamps, L. A. L., Russell, L. E., Buchholz, M. O., Bicknell, B. A., Antonov, G. K., Lau, J. Y. N., Nutbrown, R., Schmidt-Hieber, C., & Häusser, M. (2020). Targeted Activation of Hippocampal Place Cells Drives Memory-Guided Spatial Behavior. Cell, 183, 1586–1599.e1510.

Rolls, E. T. (2007). An attractor network in the hippocampus: theory and neurophysiology. Learn Mem, 14, 714–731.

Roy, A., & Narayanan, R. (2021). Spatial information transfer in hippocampal place cells depends on trial-to-trial variability, symmetry of place-field firing, and biophysical heterogeneities. Neural Netw, 142, 636–660.

Roy, R., & Narayanan, R. (2022). Ion-channel degeneracy and heterogeneities in the emergence of complex spike bursts in CA3 pyramidal neurons. J Physiol.

Roy, R., & Narayanan, R. (2023). Ion-channel degeneracy and heterogeneities in the emergence of complex spike bursts in CA3 pyramidal neurons. Journal of Physiology, 601, 3297–3328.

Royer, S., Zemelman, B. V., Losonczy, A., Kim, J., Chance, F., Magee, J. C., & Buzsaki, G. (2012). Control of timing, rate and bursts of hippocampal place cells by dendritic and somatic inhibition. Nat Neurosci, 15, 769–775.

Rubin, R., Abbott, L. F., & Sompolinsky, H. (2017). Balanced excitation and inhibition are required for high-capacity, noise-robust neuronal selectivity. Proc Natl Acad Sci U S A, 114, E9366–E9375.

Saini, S., & Narayanan, R. (2025). Degeneracy Explains Diversity in Interneuronal Regulation of Pattern Separation in Heterogeneous Dentate Gyrus Networks. Function (Oxf), 6, zqaf035.

Santhosh, A., & Narayanan, R. (2025). Diversity in the impact of heterogeneities on recurrent networks performing a cognitive task. bioRxiv, 2025.2001.2020.633872.

Seenivasan, P., & Narayanan, R. (2020). Efficient phase coding in hippocampal place cells. Physical Review Research, 2, 033393.

Sheffield, M. E., & Dombeck, D. A. (2015). Calcium transient prevalence across the dendritic arbour predicts place field properties. Nature, 517, 200–204.

Sheffield, M. E. J., Adoff, M. D., & Dombeck, D. A. (2017). Increased Prevalence of Calcium Transients across the Dendritic Arbor during Place Field Formation. Neuron, 96, 490–504 e495.

Simoncelli, E. P. (2003). Vision and the statistics of the visual environment. Current opinion in neurobiology, 13, 144–149.

Simoncelli, E. P., & Olshausen, B. A. (2001). Natural image statistics and neural representation. Annual review of neuroscience, 24, 1193–1216.

Sosa, M., Plitt, M. H., & Giocomo, L. M. (2025). A flexible hippocampal population code for experience relative to reward. Nat Neurosci, 28, 1497–1509.

Sprekeler, H. (2017). Functional consequences of inhibitory plasticity: homeostasis, the excitation-inhibition balance and beyond. Current opinion in neurobiology, 43, 198–203.

Sukenik, N., Vinogradov, O., Weinreb, E., Segal, M., Levina, A., & Moses, E. (2021). Neuronal circuits overcome imbalance in excitation and inhibition by adjusting connection numbers. Proc Natl Acad Sci U S A, 118.

Sun, Q., Jiang, Y. Q., & Lu, M. C. (2020). Topographic heterogeneity of intrinsic excitability in mouse hippocampal CA3 pyramidal neurons. J Neurophysiol, 124, 1270–1284.

Sun, Q., Sotayo, A., Cazzulino, A. S., Snyder, A. M., Denny, C. A., & Siegelbaum, S. A. (2017). Proximodistal Heterogeneity of Hippocampal CA3 Pyramidal Neuron Intrinsic Properties, Connectivity, and Reactivation during Memory Recall. Neuron, 95, 656–672 e653.

Taub, A. H., Katz, Y., & Lampl, I. (2013). Cortical balance of excitation and inhibition is regulated by the rate of synaptic activity. J Neurosci, 33, 14359–14368.

Treves, A., & Panzeri, S. (1995). The upward bias in measures of information derived from limited data samples. Neural Computation, 7, 399–407.

Van Vreeswijk, C., & Sompolinsky, H. (1996). Chaos in neuronal networks with balanced excitatory and inhibitory activity. Science, 274, 1724–1726.

Yang, J., & Prescott, S. A. (2023). Homeostatic regulation of neuronal function: importance of degeneracy and pleiotropy. Front Cell Neurosci, 17, 1184563.

Yang, W., & Sun, Q. Q. (2018). Circuit-specific and neuronal subcellular-wide E-I balance in cortical pyramidal cells. Sci Rep, 8, 3971.

Yu, L., Shen, Z., Wang, C., & Yu, Y. (2018). Efficient Coding and Energy Efficiency Are Promoted by Balanced Excitatory and Inhibitory Synaptic Currents in Neuronal Network. Front Cell Neurosci, 12, 123.

Zhou, S., & Yu, Y. (2018). Synaptic E-I Balance Underlies Efficient Neural Coding. Frontiers in Neuroscience, 12.

Zhou, S., & Yu, Y. (2018). Synaptic E-I Balance Underlies Efficient Neural Coding. Front Neurosci, 12, 46.

Zutshi, I., Valero, M., Fernandez-Ruiz, A., & Buzsaki, G. (2022). Extrinsic control and intrinsic computation in the hippocampal CA1 circuit. Neuron, 110, 658–673 e655.

